# Dissecting the KNDy hypothesis: KNDy neuron-derived kisspeptins are dispensable for puberty but essential for preserved female fertility and gonadotropin pulsatility

**DOI:** 10.1101/2022.06.07.495233

**Authors:** Inmaculada Velasco, Delphine Franssen, Silvia Daza-Dueñas, Katalin Skrapits, Szabolcs Takács, Encarnación Torres, Elvira Rodríguez-Vazquez, Miguel Ruiz-Cruz, Silvia León, Krisztina Kukoricza, Fu-Ping Zhang, Suvi Ruohonen, Diego Luque-Cordoba, Feliciano Priego-Capote, Francisco Gaytan, Francisco Ruiz-Pino, Erik Hrabovszky, Matti Poutanen, María J. Vázquez, Manuel Tena-Sempere

**Affiliations:** Instituto Maimónides de Investigación Biomédica de Córdoba (IMIBIC), Cordoba; Department of Cell Biology, Physiology and Immunology, University of Córdoba, Cordoba; Hospital Universitario Reina Sofía, Cordoba, Spain; GIGA-Neurosciences Unit. University of Liège, Liège, Belgium; Laboratory of Reproductive Neurobiology, Institute of Experimental Medicine, Budapest, Hungary; Research Centre for Integrative Physiology and Pharmacology, Institute of Biomedicine, University of Turku, Finland; Department of Analytical Chemistry, University of Córdoba, Spain; CIBER Fragilidad y Envejecimiento Saludable, Instituto de Salud Carlos III, Cordoba, Spain; CIBER Fisiopatología de la Obesidad y Nutrición, Instituto de Salud Carlos III, Cordoba, Spain

**Keywords:** Kiss1, kisspeptins, Neurokinin B, Tac2, KNDy, GnRH, gonadotropins, metabolism, puberty, fertility

## Abstract

Kiss1 neurons in the hypothalamic arcuate-nucleus (ARC) play key roles in the control of GnRH pulsatility and fertility. A fraction of ARC Kiss1 neurons, termed KNDy, co-express neurokinin B (NKB; encoded by *Tac2*). Yet, NKB- and Kiss1-only neurons are also found in the ARC, while a second major Kiss1-neuronal population is present in the rostral hypothalamus. The specific contribution of different Kiss1 neuron subsets to reproductive control remains unfolded. To tease apart the physiological roles of KNDy-born kisspeptins, conditional ablation of *Kiss1* in *Tac2*-expressing cells was implemented in vivo. Mice with *Tac2* cell-specific *Kiss1* KO (TaKKO) displayed reduced ARC kisspeptin content and *Kiss1* expression, with greater suppression in females, which was detectable at infantile-pubertal age. In contrast, *Tac2*/NKB levels were fully preserved. Despite the drop of ARC *Kiss1*/kisspeptin, pubertal timing was normal in TaKKO mice of both sexes. However, young-adult TaKKO females displayed disturbed LH pulsatility and sex steroid levels, with suppressed basal LH and pre-ovulatory LH surges, early-onset subfertility and premature ovarian insufficiency. Conversely, testicular histology and fertility were grossly conserved in TaKKO males. Ablation of *Kiss*1 in *Tac2*-cells led also to sex-dependent alterations in body composition, glucose homeostasis and locomotor activity. Our data document that KNDy-born kisspeptins are dispensable/compensable for puberty in both sexes, but required for maintenance of female gonadotropin pulsatility and fertility, as well as adult metabolic homeostasis.

**Significance Statement:** Neurons in the hypothalamic arcuate nucleus (ARC) co-expressing kisspeptins and NKB, named KNDy, have been recently suggested to play a key role in pulsatile secretion of gonadotropins, and hence reproduction. However, the relative contribution of this Kiss1 neuronal-subset, vs. ARC Kiss1-only and NKB-only neurons, as well as other Kiss1 neuronal populations, has not been assessed in physiological settings. We report here findings in a novel mouse-model with elimination of KNDy-born kisspeptins, without altering other kisspeptin compartments. Our data highlights the heterogeneity of ARC Kiss1 populations and document that, while dispensable/compensable for puberty, KNDy-born kisspeptins are required for proper gonadotropin pulsatility and fertility, specifically in females. Characterization of this functional diversity is especially relevant, considering the potential of kisspeptin-based therapies for management of human reproductive disorders.

**Disclosure Statement:** The authors have nothing to disclose in relation to the contents of this work.

## Introduction

Hypothalamic neurons producing gonadotropin-releasing hormone (GnRH) are a key hierarchical element of the neuroendocrine system governing reproduction, i.e., the hypothalamic-pituitary-gonadal (HPG) axis. These are ultimately modulated by, and integrate, multiple internal and external cues, and operate as final output pathway for the brain control of the reproductive axis (Herbison 2016). Proper pulsatile patterns of GnRH secretion are mandatory for pubertal maturation and fertility in both sexes (Herbison 2016; Maeda et al. 2010). In addition, ovulation is triggered by the so-called pre-ovulatory surge of gonadotropins, which, in turn, is driven by GnRH inputs (Maeda et al. 2010). GnRH neurons are precisely controlled by multiple regulatory signals, able to shape the pulse and surge modes of GnRH secretion (Maeda et al. 2010), and to adjust these to endogenous conditions and environmental cues. Among these afferent signals, kisspeptins, encoded by the *Kiss1* gene, have emerged in recent years as master regulators of all key aspects of reproductive maturation and function, by acting primarily on GnRH neurons, which express the canonical kisspeptin receptor, Gpr54 (aka, Kiss1r), and are potently activated by kisspeptins, in terms of electrical and secretory responses (Pinilla et al. 2012; Franssen and Tena-Sempere 2018).

In mammals, two major populations of Kiss1 neurons have been described in the hypothalamus. One is located in the arcuate nucleus (ARC), or its equivalent infundibular region in humans, while the other is found in the rostral hypothalamic area, which in rodents concentrates mainly around the anteroventral periventricular nucleus (AVPV) (Pinilla et al. 2012). These two populations participate in the stimulatory control of GnRH neurons, but are endowed with distinct roles. Thus, the ARC Kiss1 neuronal population is present in both sexes and mediates the negative feedback actions of sex steroids on the gonadotropic axis (Garcia-Galiano, Pinilla, et al. 2012). In contrast, the rostral/AVPV population of Kiss1 neurons is predominantly found in females, and it participates in mediating the positive feedback effects of estrogens and, thereby, in the generation of the pre-ovulatory surge of gonadotropins (Garcia-Galiano, Pinilla, et al. 2012). A third population of Kiss1 neurons have been reported in the amygdala (Pineda et al. 2017; Kim et al. 2011), which is also sexually dimorphic but, in contrast to the AVPV, it is more abundant in male rodents (Kim et al. 2011). The functional roles of amygdala Kiss1 neurons is still under debate, but they might contribute to the control of gonadotropin secretion and reproductive responses to environmental cues, such as odor stimuli (Aggarwal et al. 2019; Pineda et al. 2017). Additional small sets of Kiss1 neurons have been found in other brain areas, including the lateral septum, of as yet unknown functions (Yeo et al. 2016). Interestingly, projections of Kiss1 neuronal populations have been demonstrated to multiple hypothalamic and extra-hypothalamic areas (Yeo and Herbison 2011), including some not overtly related with reproductive control, therefore suggesting additional putative roles of kisspeptins in the control of other bodily functions, including metabolic homeostasis (Hudson and Kauffman 2021).

The population of Kiss1 neurons located in the ARC plays key physiological roles in the control of the reproductive axis and, hence, has attracted substantial attention in recent years. Optogenetic analyses, coupled to calcium monitoring by fiber photometry, have documented that ARC Kiss1 neurons are the core component of the hypothalamic GnRH pulse generator, which drives gonadotropin pulsatility essential for fertility (Clarkson et al. 2017). This function seemingly relies on an oscillatory circuit, composed of two main signals, neurokinin B (NKB; encoded by *Tac2* in rodents) and dynorphin (Dyn), with predominant stimulatory and inhibitory effects, respectively, on ARC Kiss1 neurons, thereby controlling the kisspeptin output onto GnRH neurons. Initial expression analyses in rodents and sheep pointed out that ARC Kiss1 neurons co-express kisspeptins, NKB and Dyn; hence, the acronym KNDy was coined to name this population (Lehman et al. 2010; Moore et al. 2018). This led to the so-called “KNDy hypothesis” for GnRH pulse generation, that proposes that KNDy neurons, which display profuse inter-connections, operate under tight auto-regulatory loops, whereby NKB stimulates, while Dyn inhibits kisspeptin release, to drive GnRH pulsatility (Moore et al. 2018). Further support for this integrative model has been provided recently in rodents, suggesting that, while KNDy neurons utilize kisspeptins, via non-synaptic appositions, to activate GnRH neurons, synchronization of KNDy neuronal network occurs in a kisspeptin-independent manner, via NKB and/or Dyn transmission (Liu et al. 2021). In addition, transient rescue of *Kiss1* expression in a fraction of *Tac2*-expressing cells in the ARC was sufficient to rescue LH pulsatility and folliculogenesis in adult female rats with whole-body congenital ablation of *Kiss1* (Nagae et al. 2021), further supporting the KNDy hypothesis for the central control of the reproductive axis.

While the KNDy hypothesis remains fully valid, compelling evidence has pointed out some degree of heterogeneity in the ARC Kiss1 neuronal population, which might depend on the species, sex and even maturational or functional stage of the reproductive axis, with evidence for the variable co-existence of KNDy, Kiss1-only and NKB-only cells in the ARC of different species (Overgaard et al. 2014; Hrabovszky et al. 2012). Thus, while ovine studies evidenced a near complete co-expression of kisspeptin, NKB and Dyn in the ARC (Moore et al. 2018), human studies have documented a much lower degree of co-localization of KNDy peptides in the infundibular region (Hrabovszky et al. 2012). Moreover, the lineage of ARC Kiss1 neurons has not been fully clarified, and previous evidence suggested that at least a fraction of this population derives from POMC precursors (Sanz et al. 2015). Collectively, the above evidence raises the possibility of specialization of different subsets of ARC Kiss1 neurons in the fine control of different facets of the functionality of the reproductive axis, especially when considered in the context of the contribution of other Kiss1 neuronal populations. However, most of the evidence on the co-expression of KNDy peptides comes from immunohistochemical and/or in situ hybridization analyses, which provide a static view of the expression status of a cell at a given moment. Hence, this approach has important limitations for interrogation of dynamic and functional aspects of the different components of the KNDy system. Thus, integral assessment of the physiological roles of KNDy-derived kisspeptins in the control of puberty, fertility and related body functions, as metabolic homeostasis, is still pending, and requires more sophisticated genetic models for precise dissection of KNDy- vs. non KNDy-born kisspeptins.

## Results

### Generation of a Tac2-cell specific Kiss1 knock-out (TaKKO) mouse line

A new mouse line, termed TaKKO (for *<u>Ta</u>c2*-specific *<u>K</u>iss1* <u>KO</u>), was generated by crossing a validated *Tac2*-Cre mouse (B6;129S-*Tac2^tm1.1(cre)Hze^*/J; JAX:021878), obtained from the Jackson Laboratories (https://mice.jax.org/), with our newly generated, Kiss1^loxP/loxP^ mouse, produced at the Turku Center for Disease Modeling (www.tcdm.fi, Turku, Finland), in which exon 3 of *Kiss1* gene is flanked by loxP sites, to allow Cre-mediated recombination. Homozygous breeding of these two lines permitted congenital ablation of exon 3 of *Kiss1*, which encodes functionally active domains of kisspeptins (Pinilla et al. 2012), in *Tac2*-cells, which prominently include ARC KNDy neurons (Lehman et al. 2010), in which *Kiss1* and *Tac2* are co-expressed (**Fig.1A**). TaKKO mice were born at a Mendelian ratio, and animals of both sexes were analyzed.

**Figure 1:**
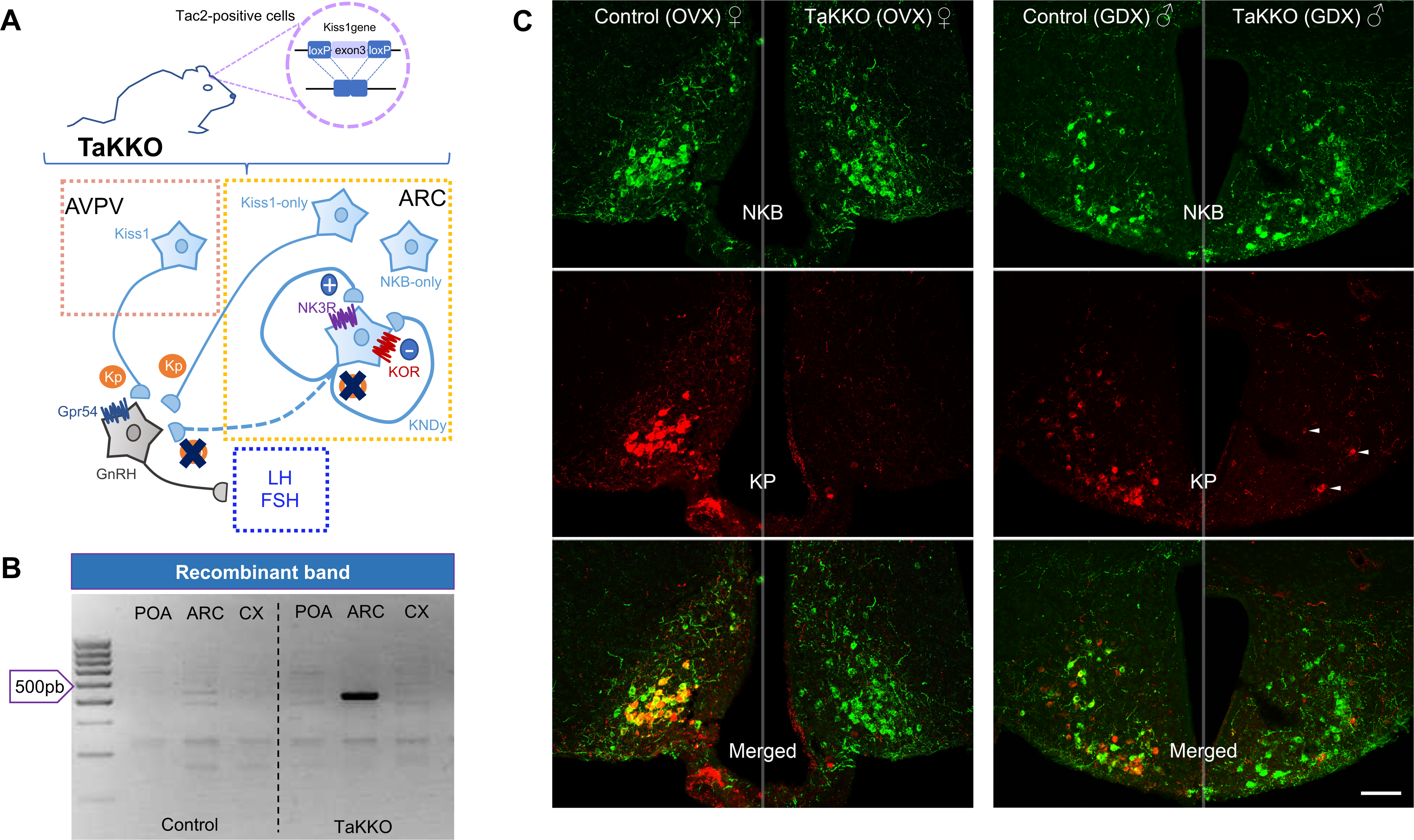
Generation of TaKKO mouse line. In **A**, schematic of generation of a novel mouse line for *Tac2*- specific *Kiss1* elimination, by crossing Tac2-Cre and Kiss1^loxP/loxP^ mouse lines. In **B**, recombinant band resulting from the deletion of *Kiss1* exon 3, detected only in the ARC of TaKKO mice. Finally, in **C**, double labelling immunohistochemistry of NKB- (green) and Kisspeptin (KP; red) in the ARC of control and TaKKO mice of both sexes. For validation analyses, mice were gonadectomized (GNX) before sampling, in order to enhance endogenous KP and NKB expression. Arrowheads point to Kiss1-only neurons in male TaKKO mice. The scale bar represents 100µm.

In line with the observed genotype, evidence for a recombinant *Kiss1* allele, after Cre-mediated excision, was obtained in TaKKO mice by PCR in the ARC, where the most abundant population of Tac2-Kiss1 (namely, KNDy) neurons is found (**Fig.1B**). In contrast, no evidence for Cre-mediated recombination was detected in the cortex or preoptic area (POA) of TaKKO mice, used as negative control, as these are devoid of substantial Tac2 expression, neither was it found in any of the tissues analyzed in control (Cre-negative) mice (**Fig.1B**). Further confirmation of the predicted phenotype was obtained by double immunohistochemistry (IHC) for detection of kisspeptin- and NKB-immunoreactivity (IR). This protocol not only documented the presence of an abundant KNDy population in the ARC of control male and female mice, which was more prominent in females, but also demonstrated the effective suppression of kisspeptin-IR in the ARC of adult TaKKO mice of both sexes, resulting in complete elimination of KNDy neurons in conditional null mice (**Fig.1C**). Yet, occasional single-labeled Kiss1-IR neurons were detected in the ARC of TaKKO mice, which were more readily detectable in the caudal portion of the ARC of males (**Fig.1C**). TaKKO mice of both sexes displayed preserved NKB-IR in the ARC, therefore supporting lack of ablation of NKB in our *Tac2*-driven Kiss1 null mouse. Since this proof-of-principle study intended to be mainly qualitative, gonadectomized mice of control and TaKKO genotypes were used, as a means to enhance endogenous *Kiss1*/kisspeptin expression (Pinilla et al. 2012), to ease detection of any kisspeptin-expressing neurons in TaKKO mice.

### Changes in hypothalamic Kiss1 expression in TaKKO male and female mice during postnatal maturation

The profiles of *Kiss1* expression at the two main hypothalamic sites, i.e., ARC and AVPV, were assessed in TaKKO mice of both sexes, using quantitative in situ hybridization (ISH), at different stages of postnatal maturation, i.e., infantile (2-wks of age; corresponding to mini-puberty), peri-pubertal (4-wks) and young adult (2-months). The number of *Kiss1*-expressing cells was significantly suppressed in the ARC of TaKKO female mice. A drop of >85% that was detectable already at the infantile period and persisted up to adulthood (**Fig.2A-B**). In contrast, *Kiss1* expression was unaltered in the ARC of TaKKO male mice, at 2-weeks and 4-weeks of age (albeit a trend for decrease was observed at the latter age-point), whereas a significant (∼40%) decrease in the number of ARC *Kiss1*-expressing cells was found in TaKKO male mice at adulthood (**Fig.2C-D**).

**Figure 2:**
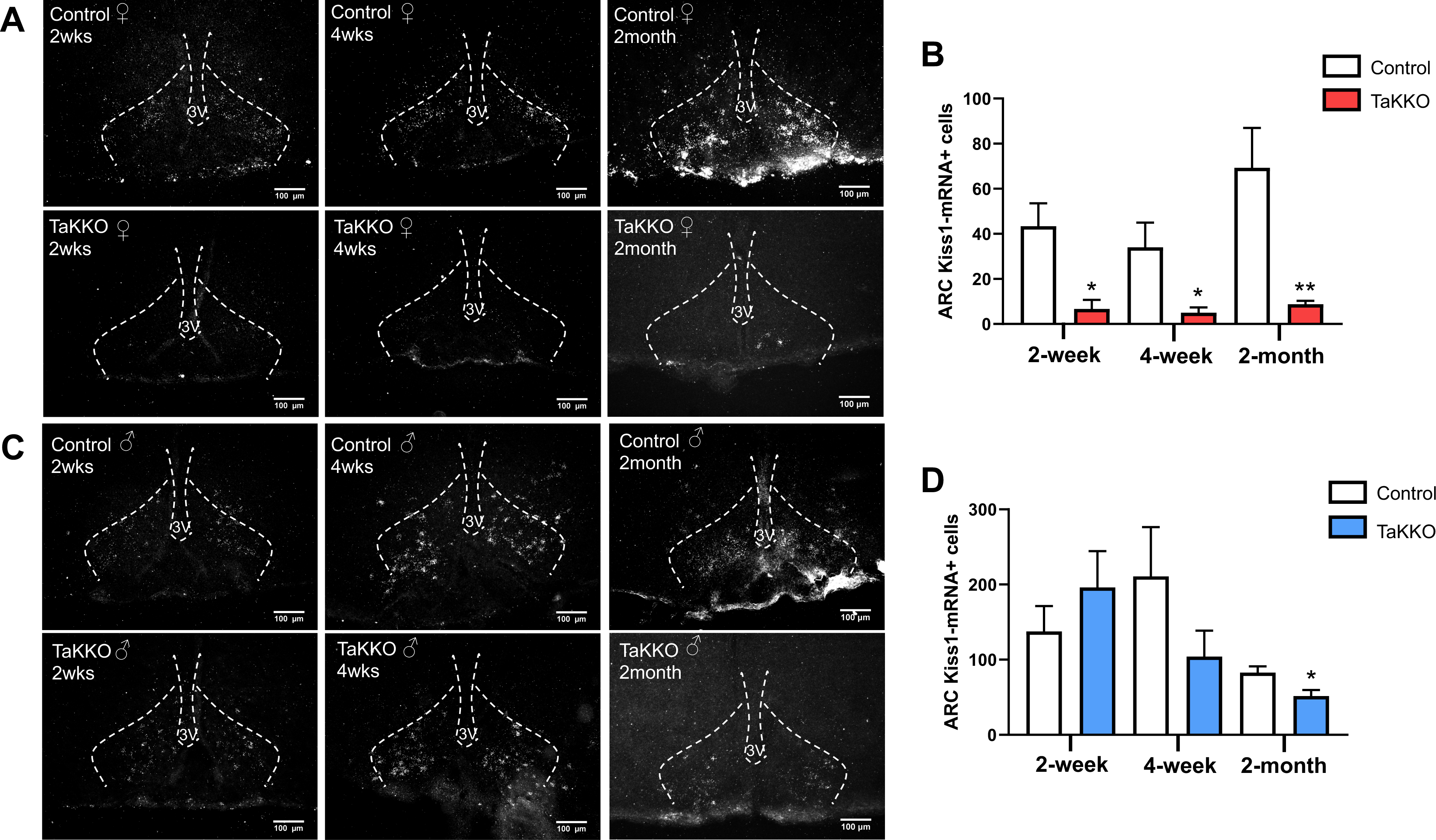
Changes in Kiss1 expression in the ARC of TaKKO mice along postnatal maturation. In **A**, representative in-situ hybridization (ISH) photomicrographs are shown of *Kiss1*-expressing neurons in the anterior region of the ARC from 2-week (infantile period, coinciding with mini-puberty), 4-week (peri-puberty) and 2-month-old (young-adult) control (white-bars) and TaKKO female (red-bars) mice; Controls: n= 5, 6 and 4; TaKKO: n= 3, 5 and 5, at 2-wks, 4-wks, and 2-mo, respectively. Quantification of *Kiss1* expression in female mice at different ages is displayed in **B**. In **C**, representative ISH photomicrographs are presented of *Kiss1*-expressing neurons in the anterior region of the ARC from 2-wks, 4-wks and 2-month-old control (white-bars) and TaKKO male (blue-bars) mice; Control: n=5, 5 and 4; TaKKO n= 5, 5, and 4 at 2-wks, 4-wks and 2-mo, respectively. Quantification of *Kiss1* expression is displayed in **D**. Data are presented as mean ± SEM. Statistical significance was determined at each age-point by Student t-test: *P<0.05 and **P<0.01 vs. corresponding control group at different ages. The scale bar represents 100µm.

Similar *Kiss1* ISH analyses were conducted in the AVPV of TaKKO mice; yet, negligible expression at the infantile period prevented us from quantification at this period. Kiss1-expressing cells in female mice were much more abundant than in males of both genotypes, at peri-pubertal and adult periods. Yet, while in 4-week-old mice, no significant differences were found between genotypes, in young adult (2-mo-old) mice, *Kiss1* mRNA expression was significantly decreased in the AVPV of TaKKO female mice (**Suppl.Fig-S1**), whereas an opposite, non-significant trend was detected for TaKKO males. However, the significant drop of AVPV *Kiss1* expression detected in adult TaKKO females was fully prevented by estradiol supplementation in ovariectomized TaKKO mice (**Suppl.Fig-S1**). This suggests that this drop is not primary caused by *Tac2*-driven Kiss1 recombination, but rather secondary to the decline in ovarian function presumably linked to the dramatic reduction of ARC *Kiss1* expression in TaKKO female mice, since reduction of sex steroid levels is known to lower Kiss1 expression in the AVPV (Garcia-Galiano, Pinilla, et al. 2012).

Finally, despite the marked decline of ARC *Kiss1* mRNA levels in TaKKO mice, no significant alteration in *Tac2* (encoding NKB) expression was found in the ARC of young adult TaKKO animals, in either females or males. These findings confirm our initial IHC data and supports a compartmentalized ablation of kisspeptins in KNDy neurons, without affecting other KNDy transmitters, as NKB (**Suppl.Fig-S2**).

### Sex-biased impact of conditional ablation of Kiss1 in KNDy neurons upon reproductive parameters

Comprehensive reproductive characterization was conducted in TaKKO mice of both sexes. First, control and TaKKO male and female mice were examined for phenotypic signs of puberty onset. Despite the strong decrease of ARC *Kiss1* mRNA at the infantile and peripubertal period, TaKKO females displayed normal body weight gain (**Suppl.Fig-S3A**), as well as complete vaginal opening (VO; **Suppl.Fig-S3B**) and first estrus, as marker of first ovulation (**Suppl.Fig-S3C**), at similar ages than control females. In addition, ovarian histological analysis did not detect clear differences between peri-pubertal control and TaKKO female mice, and in both groups, the ovaries contained high numbers of growing follicles, with the largest healthy follicles being early/mid antral follicles (**Suppl.Fig-S3F**). Likewise, TaKKO males failed to display significant differences in body weight gain after weaning or in the age of preputial separation, as external sign of male puberty (**Suppl.Fig-S3D,E**).

In contrast, conditional ablation of *Kiss1* in *Tac2*-expressing cells evoked marked alterations of reproductive parameters in young adult mice, which occurred in a sexually-divergent manner. Thus, while TaKKO female displayed significantly suppressed basal LH levels, without changes in FSH concentrations, as well as reduced ovarian and uterus weights, adult TaKKO males did not show changes in basal gonadotropin levels or testicular weight (**Suppl.Fig-S4**). In addition, conditional ablation of *Kiss1* expression in *Tac2* cells totally prevented the rise of serum LH levels following elimination of the negative feedback from sex steroids by gonadectomy (GNX) in females, whereas the post-GNX LH response was partially blunted but not totally ablated in adult TaKKO males (**Suppl.Fig-S4**). Sex steroid profiling in TaKKO mice of both sexes, conducted at 4-weeks and 2-months of age, allowed further characterization of the hormonal profiles. In females, the increase in serum levels of progesterone (P) detected between pubertal and young adult control mice was abrogated in TaKKO animals, whose P levels in adulthood tended to be lower than in controls (P=0.09). A similar trend for decreased serum levels was detected for testosterone (T) and estradiol (E_2_) in adult TaKKO females. In the case of males, no differences in basal levels of testosterone or DHT were found between TaKKO and controls at the two age-points; yet, a significant decrease of progesterone levels was detected in adult TaKKO males (**Suppl.Fig-S5**). Altogether, these data pointed to a more prominent impact of conditional Kiss1 ablation in females, that were accordingly submitted to further characterization.

**Figure 4:**
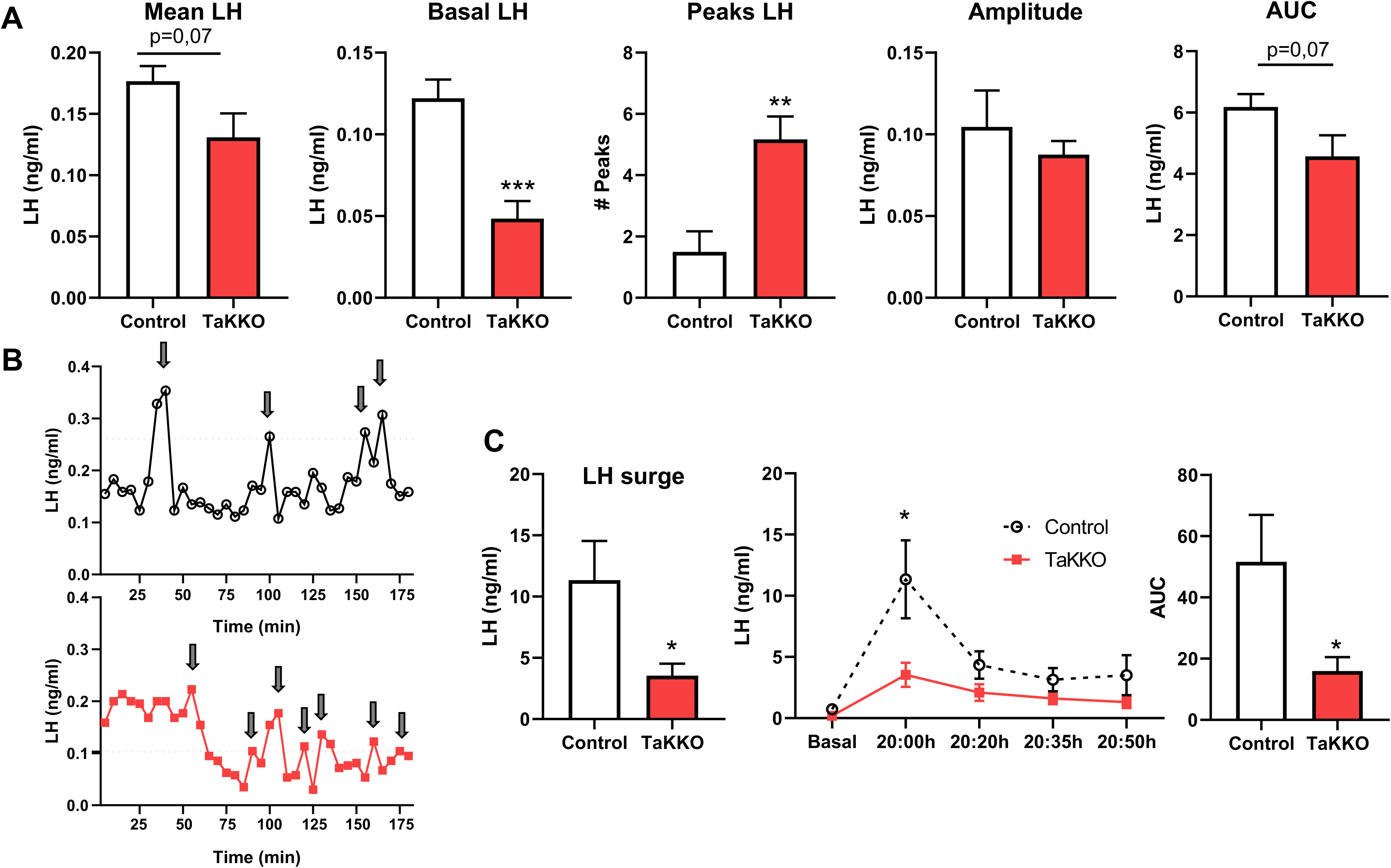
Fertility markers of young-adult TaKKO female mice. In **A**, indices of estrous cyclicity, including frequency of cycles (# of complete cycles over a 10-day-period) and representative cyclic profiles, are shown from control (black-symbols) and TaKKO (red-symbols) female mice. In **B**, ovulatory efficiency, denoted by number of corpora lutea per ovary, and representative histological images of ovarian sections from control and TaKKO female mice (Control n=5; TaKKO n=5) are presented. In controls, two generations of corpora lutea were found indicating normal cyclicity; in contrast, most TaKKO females showed only one generation of corporal lutea, either fresh or regressing, indicating sporadic or defective ovulation. In **C**, rates of successful pregnancies and average litter sizes of control and TaKKO females mated with control males of proven fertility, are displayed (Control n=5; TaKKO n=5). Similar parameters are shown in **D** from control and TaKKO males, mated with control females (Control n=5; TaKKO n=6). CL: Corpus luteum; rCL: regressing CL.

**Figure 5:**
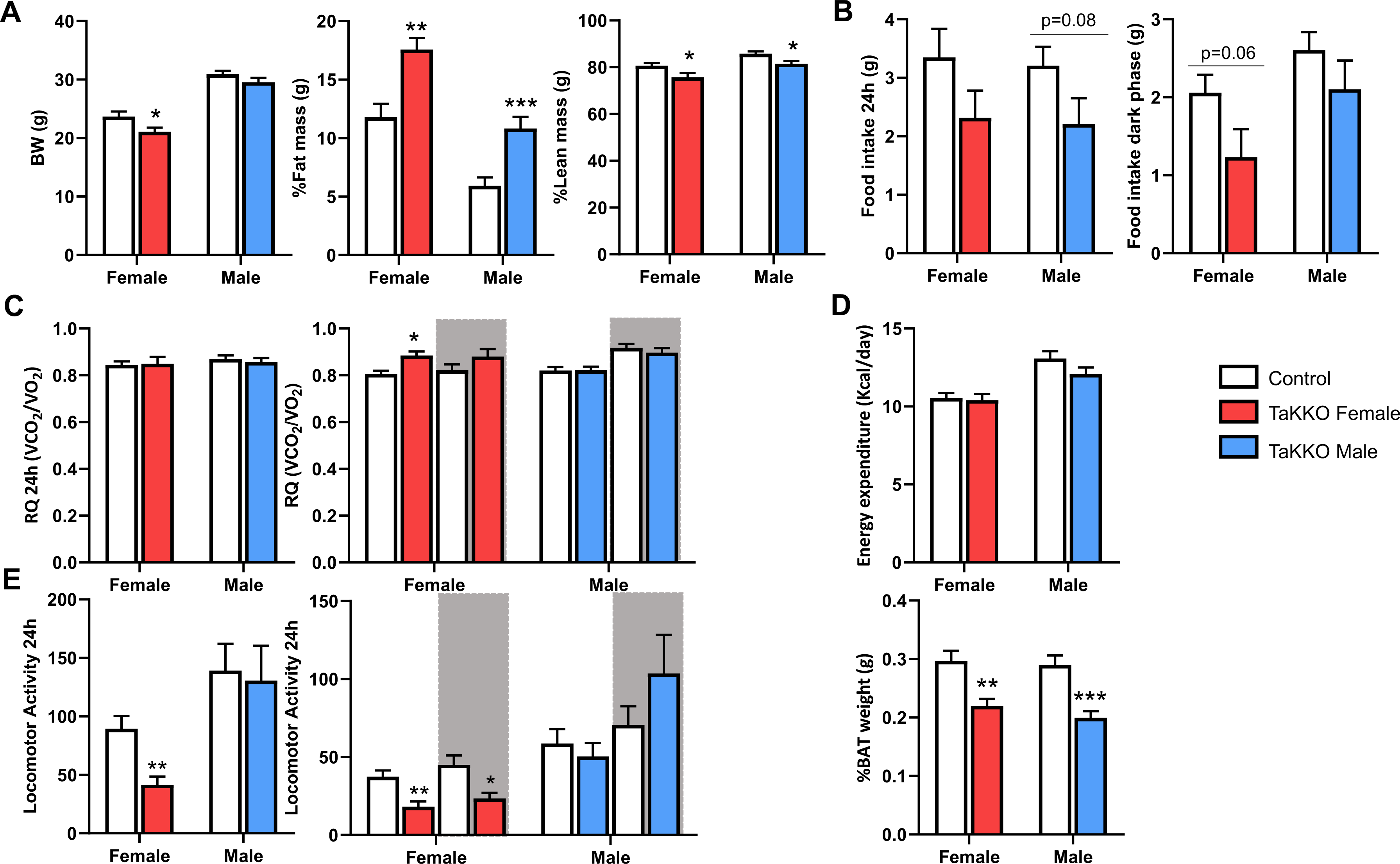
Basic metabolic characterization of TaKKO mice of both sexes. In **A**, body-weight (BW) and body-composition (percentage of fat mass and lean mass) are presented from control and TaKKO mice (females: Control n=12, TaKKO n=8; males: Control n=13, TaKKO n=10). Additionally, spontaneous feeding patterns, defined by total food intake, in a 24h-period and during the dark phase, are presented in **B** (females: Control n=8, TaKKO n=6; males: Control n=8, TaKKO n=7). In **C**, respiratory quotient (RQ), either measured in a 24h-period or split between light- and dark-phases, is shown (females: Control n=10 and TaKKO n=7; males: Control n=9 and TaKKO n=9). In **D**, energy expenditure, measured by indirect calorimetry over a 24h-period, as well as relative weights of brown adipose tissue (BAT), are shown from control and TaKKO mice (female: Control n=10 and TaKKO n=7; male: Control n=8 and TaKKO n=8). Finally, in **E**, total locomotor activity, in a 24h-period and split between light- and dark-phases, is presented from control and TaKKO mice (female: Control n= 8, TaKKO n=6; male: Control n= 9, TaKKO n=9). Data are shown as mean ± SEM. Statistical significance of differences between genotypes was determined by Student t-test for each sex: *P<0.05; **P<0.01 and ***P<0.001 vs controls.

In line with our initial single-point determinations, analyses of pulsatile LH secretion of young adult TaKKO females revealed major alterations in LH secretory patterns. Thus, assessment of the profile of pulsatile LH release during a 3-h period in TaKKO females at diestrus demonstrated a nearly-significant lowering of mean LH levels (P=0.07) and a significant drop of basal LH level (**Fig.3A**), which appeared to be partially offset by an increase in the number of LH pulses (**Fig.3A,B**). Yet, no significant differences in terms of pulse amplitude were detected in TaKKO females, whereas the apparent decline in integral LH secretory mass over the 3-h period, calculated as area under the curve (AUC), did not reach statistical significance (P=0.07). In addition, the capacity of TaKKO female mice to display estrogen-primed LH secretory peaks, reminiscent of the pre-ovulatory surge, was explored, using a validated model of E_2_ supplementation in ovariectomized (OVX) animals. Our protocol of E_2_ priming induced a robust LH secretory surge in control females, while this was markedly attenuated in TaKKO mice, as denoted by a significant decrease of LH concentration at the peak, and substantially suppressed LH release profile and AUC secretory mass (**Fig.3C**). These data suggest that both the tonic and pulse modes of LH secretion are altered in TaKKO female mice.

**Figure 3:**
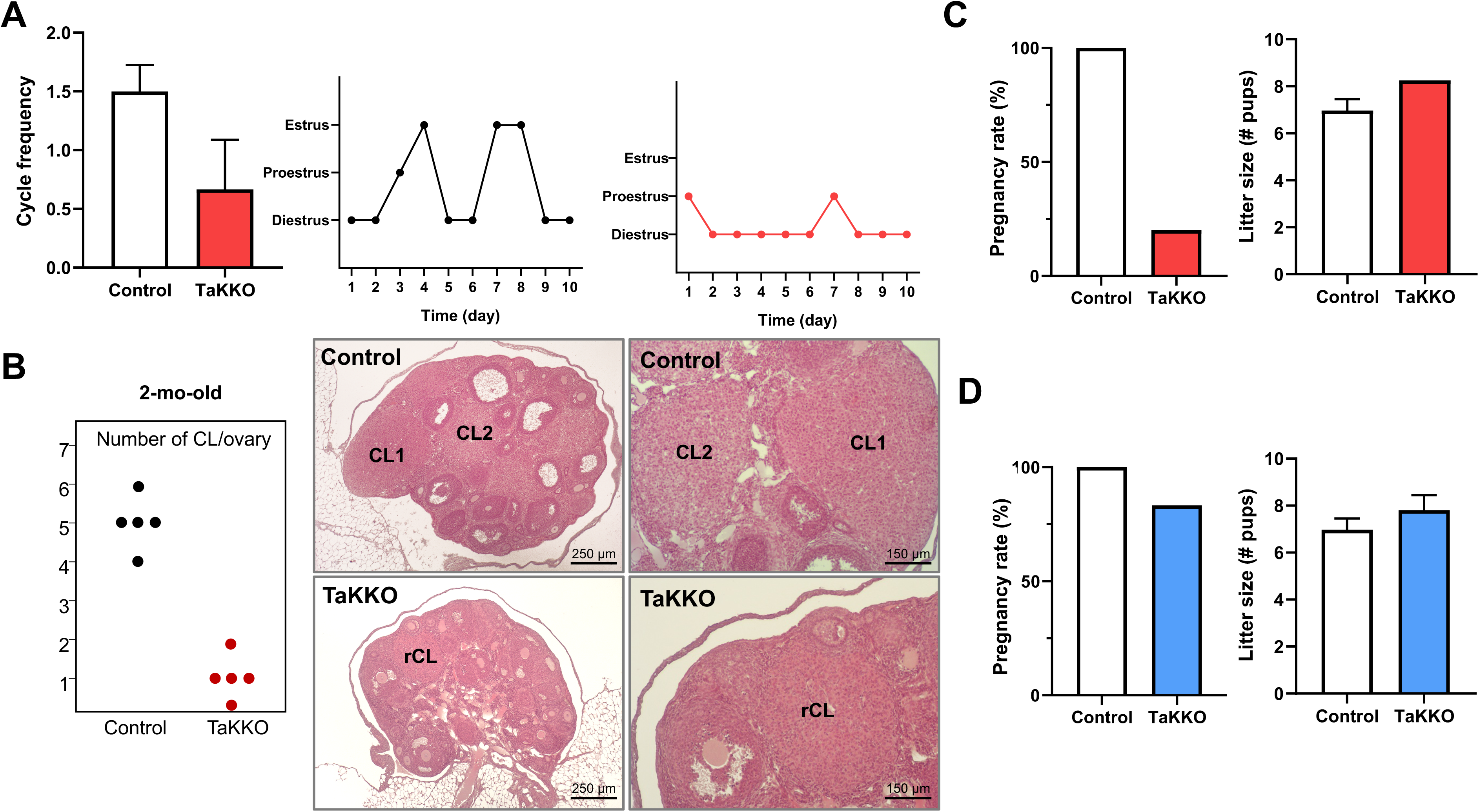
LH pulsatile secretion and LH surges in TaKKO female mice. In **A**, mean and basal LH levels, as well as number of LH pulses, pulse amplitude and area-under-the-curve (AUC) of total pulses, over a 3-hour sampling period, are represented from control and TaKKO female mice (Control n=6; TaKKO n=6). In **B**, individual representative LH secretory profiles of control (black-line) and TaKKO (red-line) female mice are shown. Finally, in **C**, different parameters of estrogen-primed, LH surges in control and TaKKO female mice (Control n=6; TaKKO n=6), ovariectomized and supplemented with E_2_, are presented; these include peak LH levels, time-course profiles and global AUC values of LH surges. Data are presented as mean ± SEM. Statistical significance was determined by Student t-test: *P<0.05, **P<0.01 and ***P<0.001 vs. controls. Black arrows represent LH pulses.

In line with these alterations, estrous cyclicity, as assessed by daily vaginal cytology, was perturbed in young adult TaKKO females (**Fig.4A**), which displayed also clear morphometric changes in the ovary. Thus, while control females were cycling, with abundant growing follicles at different maturational stages, and, at least, two generations of corpora lutea, TaKKO females displayed variable ovarian phenotypes. These ranged from cyclic mice with two generations of corpora lutea, to occasional mice lacking corpora lutea (i.e., non-ovulating) or, predominantly, female mice with only one corpus luteum generation, either fresh or regressing, suggesting sporadic ovulation. In all cases, small growing and antral follicles were abundant. Notably, in TaKKO mice with preserved ovulation, the number of corpora lutea of the current cycle was lower than in controls, as sign of decreased ovulatory capacity (**Fig.4B**). In good agreement, fertility tests denoted a dramatic drop in fecundity of TaKKO female mice, with an 80% drop in birth rates, without apparent changes in litter size (**Fig.4C**). In contrast, adult TaKKO male mice displayed largely preserved fertility, with birth-rates and litter-sizes similar than controls (**Fig.4D**). In addition, testicular histology, including spermatogenesis and interstitial Leydig cells, was normal in TaKKO males, even up to 12-months of age (*data not shown*).

LH secretory profiles and ovarian histology were also studied in older (12-month-old) female mice of both genotypes. These analyses further documented the impact of conditional ablation of *Kiss1* in *Tac2*-cells, which was aggravated with age, as denoted by a significant reduction of basal LH levels and an increase in the number of LH pulses. In addition, aged TaKKO females showed no regular estrous cycles, staying mainly in diestrus, as compared to age-matched controls that still displayed normal estrous cyclicity. Alike, ovarian histology in aged TaKKO females revealed the absence of corpora lutea, as sing of ovulatory failure, although small growing follicles were abundant, while healthy follicles were scarce and corresponded to early antral follicles. In contrast, pair-aged controls showed normal ovarian phenotypes and contained at least two generations of corpora lutea, as index of maintained cyclicity (**Suppl.Fig-S6**).

### Metabolic impact of conditional ablation of Kiss1 in KNDy neurons in both sexes

Given the proposed roles of kisspeptins in metabolic control, TaKKO mice of both sexes were subjected also to metabolic phenotyping. Male and female TaKKO mice displayed a significant increase of adiposity (denoted by % fat mass), together with significant, albeit moderate reduction of lean mass, as compared to controls. In addition, a modest decrease of body weight (BW) was observed in TaKKO females, whereas TaKKO males showed no changes in BW (**Fig.5A**). These alterations in body composition were not linked to significant changes in food intake, which was not altered in TaKKO animals, although a trend for decrease food intake was detected in both sexes, either during the 24h-period or the dark-phase (**Fig.5B**). In addition, the respiratory quotient (RQ) was studied during a period of 24 hours, and light vs. dark phases were distinguished. Only TaKKO females showed a significant alteration, with increased RQ during light-phase, suggesting increased utilization of carbohydrates (**Fig.5C**). Moreover, while indirect calorimetry over a 24h-period revealed no differences in energy expenditure in TaKKO mice of both sexes (**Fig.5D**), relative weight of brown adipose tissue was significantly decreased in male and female TaKKO mice (**Fig.5D**). Additionally, locomotor activity was significantly decreased during light and dark phases in TaKKO females compared to control females (**Fig.5E**), while no significant changes were observed in TaKKO males (**Fig.5E**).

Further metabolic characterization included glucose homeostasis and body temperature analyses. Marginal changes in glucose and insulin tolerance were detected in TaKKO mice, especially in males (**Suppl.Fig-S7**); despite enhanced adiposity, GTT and ITT were improved in conditional null animals. In detail, peak glucose levels at GTT were lower in TaKKO males at 20-min after the glucose bolus, while they showed enhanced insulin sensitivity, as denoted by ITT. In females, glucose tolerance was marginally enhanced, with lower glucose levels at 60-min after glucose bolus, while insulin sensitivity was similar in control and TaKKO females. In addition, body temperature was analysed also in TaKKO mice. Tail temperature was measured every 10-min for 24-hours in female mice; yet, no significant differences were noted in maximal and minimum temperatures, neither was the global decrease in tail temperature during the dark phase affected in TaKKO females. In addition, no differences in inter-scapular temperature, as surrogate marker of brown adipose activity, were detected between TaKKO and control male and female mice, at either light- or dark-phases (**Suppl.Fig-S8**).

## Discussion

In recent years, KNDy neurons have been recognized as key component of the GnRH pulse generator and, hence, major hub for the precise control of the reproductive axis. This neuronal population expresses indispensable signals for proper reproductive maturation and function, as kisspeptins and NKB (Seminara et al. 2003; Topaloglu et al. 2009; Topaloglu et al. 2012). In addition, neuroanatomical data pointed out the existence of auto-regulatory loops whereby NKB and Dyn might reciprocally control the output of kisspeptins onto GnRH neurons (Moore et al. 2018). All these data led to the proposal of the KNDy hypothesis for the control of GnRH pulsatility. While this hypothesis remains fully valid, it tended to uniform all ARC Kiss1 neurons as a single functional unit. However, fragmentary evidence suggests that, depending on the species, sex and lineage, the population of Kiss1 neurons in the ARC is heterogeneous; a feature whose functional relevance remains largely neglected. Here, we report the generation and comprehensive characterization of a novel mouse model, with congenital ablation of *Kiss1* in *Tac2*- expressing cells, as platform for the integral analysis of the specific roles and relative importance of KNDy-born kisspeptins in the physiological control of reproductive maturation and function, as well as other related bodily systems, such as metabolic homeostasis. This model allows us also to conduct detailed sex comparisons in a physiological setting, since key aspects of neurohormonal regulation in general (Miguel-Aliaga 2022), and Kiss1 neurons in particular, are subjected to important sex differences.

While evidence for the heterogeneity of the ARC population of Kiss1/KNDy neurons had been provided by a number of expression studies in different species (Overgaard et al. 2014; Hrabovszky et al. 2012), assessment of the actual proportion of Kisspeptin/NKB co-expression remains inconclusive and has been based so far on neuroanatomical data, which provides kind of static snapshot of co-localization of these KNDy peptides at a given age and functional state of the HPG axis. Conversely, our genetic model enables us to dynamically assess the percentage of neurons co-expressing kisspeptins and NKB along maturation, as Cre-mediated *Kiss1* ablation would only occur in cells expressing *Tac2* at any point of their lifetime.

Our data not only conclusively document the existence of Kiss1-only cells (i.e., not co-expressing the other key KNDy peptide, NKB, at any point during maturation) in the ARC in both males and females, but are also the first to unambiguously substantiate a marked sex difference in the proportion of *Kiss1*/*Tac2* expressing cells, which is much larger in females. Thus, the dramatic drop of *Kiss1* mRNA in the ARC of TaKKO female mice reveals that <15% of *Kiss1* expression comes from non-KNDy cells, whereas in males this proportion rises to >60%, based on expression data in young adult animals. These findings are in line with our previous cross-sectional immunohistochemical studies suggesting that neurons co-expressing NKB and kisspeptins are more prominent in the caudal ARC of adult female rats, while neurons expressing NKB, but not kisspeptins, are more abundant in the ARC of male rats (Overgaard et al. 2014). Altogether, these data suggest a greater role of KNDy-derived kisspeptins in the control of female reproduction, as supported also by our functional studies. Moreover, our time-course analyses in infantile and peri-pubertal mice disclosed striking sex-differences in the developmental program of co-expression of *Tac2* and *Kiss1* in the ARC during postnatal maturation. While a marked suppression of *Kiss1* was detectable already at PND15 in TaKKO females, in males, such co-expression seemingly starts later, with a significant decrease in *Kiss1* mRNA levels in the ARC being observed in TaKKO males only after puberty. However, completion of pubertal maturation was observed not only in males but also in TaKKO females, which attained first ovulation. This is in clear contrast with the reported failure of sexual maturation of Gpr54 and Kiss1 null female mice (Seminara et al. 2003; d’Anglemont de Tassigny et al. 2007; Garcia-Galiano, van Ingen Schenau, et al. 2012), globally devoid of kisspeptin signaling congenitally, and suggests that KNDy-born kisspeptins are dispensable (or compensable) for female puberty onset, which could be driven by kisspeptin inputs from non-*Tac2* cells, either in the ARC or elsewhere, in our TaKKO model. Of note, evidence for substantial redundancy and/or compensation of kisspeptin input as pubertal driver had been previously proposed, using models of congenital suppression of *Kiss1* or Kiss1 neuronal ablation (Popa et al. 2013; Mayer and Boehm 2011). Yet, those genetic studies evoked an indiscriminate reduction of *Kiss1* expression (>95%), and did not account for potential divergences between different sources of kisspeptin input. Our current findings refine these observations, supporting that safeguard mechanisms operate to maintain female pubertal progression, despite suppression of KNDy-born kisspeptins at mini-puberty (i.e., PND15).

While puberty onset was preserved in TaKKO mice of both sexes, divergent, sex-dependent trajectories in terms of functionality of the reproductive axis were detected in post-pubertal female vs. male mice lacking *Kiss1* expression in *Tac2*-cells, which were congruent with the differential impact in terms of ARC *Kiss1* expression. Thus, key reproductive parameters, such as basal LH and FSH levels, as well as testicular weight and microstructure, were normal in TaKKO males, which were grossly fertile and displayed fully conserved testicular histology even at 1-year of age. In clear contrast, TaKKO females displayed phenotypic features of premature reproductive impairment, that became detectable even at 2-months of age. In fact, young-adult TaKKO females not only showed lower basal LH levels and decreased ovarian and uterus weights, but presented also clear signs of ovulatory insufficiency, denoted by estrous cycle irregularity, markedly decreased numbers of corpora lutea and suppressed fecundity at this early age. These features occurred despite fully preserved *Tac2* expression, suggesting conserved NKB input, and were aggravated at 1-year of age, when all TaKKO female were anovulatory, in contrast with the preserved cyclicity of pair-aged control female mice. These data fully support the greater role of KNDy- born kisspeptins in the control of female reproduction, and evidence that the non-KNDy kisspeptin component, although able to safeguard pubertal progression, is insufficient to maintain ovarian function in adulthood. The fact that markers of incipient ovulatory failure were detectable already at 2-months, which progressed into ovulatory arrest before 12-months of age, is reminiscent of a state of premature ovarian insufficiency (POI), which may recapitulate some phenotypic features of late-onset forms of POI, which are the most common in humans (Beck-Peccoz and Persani 2006).

Compelling data gleaned in recent years have documented that KNDy neurons in the ARC are a central component of the GnRH pulse generator (Clarkson et al. 2017). Using our TaKKO model, coupled to serial, ultra-sensitive LH determinations, we have further interrogated this regulatory system in a physiological setting. Conditional ablation of kisspeptins from KNDy cells did not prevent pulsatile LH secretion, but substantially disrupted LH secretory profiles in young adult TaKKO female mice. Major changes included a significant lowering of basal LH levels, together with a rise of the number of LH pulses, while total LH secretory mass and mean LH levels tended to be (non-significantly) decreased. Whereas these changes attest for a role of KNDy-born kisspeptins in the definition of GnRH secretory patterns, they also help to dissect out the contribution of KNDy- vs. non-KNDy kisspeptins in the regulation of different components of the pulsatile secretion of GnRH. Thus, preserved KNDy-born kisspeptin input is essential for keeping basal LH levels and possibly LH secretory mass, but is dispensable for actual pulse generation, at least in our genetic model. The most tenable explanation is that non-KNDy kisspeptins (e.g., from Kiss1-only neurons) can overcome/compensate the lack of kisspeptin output from *Tac2*-cells to drive GnRH pulses. In fact, the increased number of LH pulses in TaKKO female mice possibly reflects a compensatory mechanism to maintain GnRH neurosecretion despite conditional ablation of KNDy-born kisspeptins. Preserved kisspeptin output from Kiss1 neurons outside ARC and/or conserved ARC NKB signals may contribute also to this phenomenon. The fact that similar changes of LH secretory profiles were observed in 1-year-old TaKKO female mice, with significantly suppressed basal LH levels but increased LH pulse frequency, further corroborates the consistency of these findings. Recent data suggest that normal LH pulsatility can be maintained by rescue of >20% of KNDy neurons in global Kiss1 null female rats, while >90% suppression of ARC Kiss1 expression was sufficient to abolish LH pulses (Nagae et al. 2021). These studies, however, did not recognize the potential diversity across Kiss1 neurons in the ARC. Our present data showing decreased basal LH levels but increased LH pulsatility in female mice with conditional ablation of *Kiss1* selectively in KNDy cells complement these previous findings and conclusive document preserved GnRH pulse generation capacity after selective ablation of KNDy-born kisspeptins.

Due to the lack of changes of basal gonadotropin levels and overt alterations in other reproductive indices, LH pulsatility was not assessed in TaKKO males. Yet, removal of negative feedback by GNX surfaced that, under extreme conditions, the male gonadotropic axis displays also functional alterations in TaKKO mice. Thus, the post-GNX rise of LH levels was not only totally prevented in TaKKO females, but it was also partially compromised in male mice with conditional elimination of KNDy-born kisspeptins. Since the rise of *Kiss1* expression in the ARC is known to be determinant for the elevation of LH following GNX in both sexes (Garcia-Galiano, Pinilla, et al. 2012), our data in males suggest that, although non- KNDy kisspeptins are sufficient to maintain LH secretion in basal conditions, the KNDy kisspeptin component is mandatory for full LH hyper-secretion after removal of gonadal sex steroids, which are known to play a fundamental role in the modulation of the tonic secretion of gonadotropins also in the male (Tsukamura 2022).

Besides tonic secretion, in females, gonadotropins are also released at the pre-ovulatory period under a surge mode, leading to the pre-ovulatory peak that drives ovulation (Tsukamura 2022). Based on differential responses to estrogens, which suppress *Kiss1* expression in the ARC but increase *Kiss1* levels in the AVPV, the latter population of Kiss1 neurons has been proposed as major conduit for the positive feedback of estrogens, that triggers the pre-ovulatory surge of gonadotropins (Sobrino et al. 2022; Wang and Moenter 2020). In contrast, ARC Kiss1 neurons have been considered marginal in this phenomenon, even if cellular ablation experiments documented that elimination of KNDy neurons in the ARC enhanced the pre-ovulatory surge of LH (Mittelman-Smith et al. 2016). Our sex steroid-priming experiments demonstrate that KNDy-derived kisspeptins not only contribute to the tonic control of LH secretion, but modulate also the surge mode of LH secretion, as the magnitude of estrogen-primed LH secretion was significantly blunted in TaKKO females. To our knowledge, this is the first functional evidence for a role of ARC KNDy-born kisspeptins in the positive regulation of the pre-ovulatory surge, as ablation of such KNDy-component of kisspeptin input resulted in significantly suppressed LH surges. While the precise circuit for such phenomenon is yet to be disclosed, the existence of projections of ARC Kiss1 neurons to the rostral preoptic hypothalamic area provides the anatomical substrate for a putative KNDy-born kisspeptin output to the AVPV (Yeo and Herbison 2011). Of note, comparison of the impact of KNDy neuronal ablation (Mittelman-Smith et al. 2016) vs. selective elimination of KNDy-born kisspeptins, which evoked enhancement or suppression of the magnitude of the LH surge, respectively, surfaces a previously unsuspected dual control of the surge mechanisms by ARC KNDy neurons, with stimulatory and inhibitory signals. While the nature of such inhibitory signals has not been elucidated and can only be speculated (e.g., whether Dyn may also inhibit AVPV Kiss1 neurons, as proposed in the ARC), our current data demonstrate that kisspeptins from KNDy neurons seemingly contribute to the stimulatory control of the pre-ovulatory surge of LH driving ovulation. The possibility that the defective LH surge seen in our TaKKO female mice may stem from the congenital ablation of at least part of kisspeptin production in the AVPV can be ruled out, since TaKKO females under physiological estradiol replacement displayed fully preserved *Kiss1* levels in this rostral hypothalamic area. Furthermore, recent RNA-seq analyses for identification of actively translated mRNA transcripts in AVPV Kiss1 neurons failed to detect *Tac2* expression in this neuronal population, despite the observed expression of other KNDy genes, such as *Kiss1* and *Pdyn* (Stephens and Kauffman 2021). This feature further validates our approach of *Tac2*- driven Cre expression for conditional ablation of *Kiss1* in ARC KNDy, but not in AVPV Kiss1 neurons.

Genetic models have been instrumental to demonstrate the key physiological roles of kisspeptins and their receptor, Gpr54, in the control of gonadotropin secretion and the reproductive axis (Colledge et al. 2013), including initial assessment of the main site of action of kisspeptins (Kirilov et al. 2013), and major mechanisms of Kiss1 regulation, e.g., its control by estrogen receptors and leptin (Donato et al. 2011; Mayer et al. 2010). Most of these models, however, failed to discriminate between different populations of Kiss1 neurons or sources of kisspeptin input to GnRH neurons, and thus fell short in capturing the potential heterogeneity of KNDy neurons. Our model of conditional ablation of *Kiss1* in *Tac2*-expressing cells partially circumvents such limitation, as it allows not only selective targeting of ARC Kiss1 expression, but also in vivo dissection of the roles of KNDy- vs. non KNDy-derived kisspeptins within the ARC in the control of different facets of the HPG axis. Very recently, characterization of a mouse line with conditional ablation of *Kiss1* in *Pdyn*-expressing cells has been reported (Nandankar et al. 2021). Notably, despite evidence for limited co-expression of *PDYN* and *KISS1* genes in the infundibular population (equivalent to ARC) of Kiss1 neurons in humans (Hrabovszky et al. 2012), this approach resulted in virtually complete ablation of *Kiss1* expression in the ARC in mice, so that this Pdyn-Cre/Kiss1^fl/fl^ line was considered as model of ARC-specific ablation of kisspeptins, which largely recapitulated the phenotypic features of other models of indiscriminate ARC-directed suppression of Kiss1 expression (Padilla et al. 2019; Nagae et al. 2021). Considering the proposed co-expression of *Kiss1*, *Tac2* and *Pdyn* in discrete neuronal populations in the ARC, comparison of our phenotypic findings with those of the Pdyn-Cre/Kiss1^fl/fl^ mouse further documents the heterogeneity of KNDy neuronal populations (**Suppl. Table S1**). Thus, while both TaKKO and Pdyn-Cre/Kiss1^fl/fl^ lines displayed grossly preserved pubertal maturation, the impact of *Pdyn*-mediated ablation of *Kiss1* was larger than that of *Tac2*-induced Kiss1 elimination in adult mice of both sexes. This was especially evident in males, since Pdyn-Cre/Kiss1^fl/fl^ mice displayed suppressed pulsatile LH secretion, testicular histological alterations and subfertility, while adult TaKKO males did not show overt signs of reproductive impairment, with conserved spermatogenesis up to one-year of age. Similarly, whereas overall fertility and basal LH levels were severely compromised already in our young-adult TaKKO females, alteration of other parameters, such as the number of LH pulses and serum estradiol levels, was much milder than in adult Pdyn-Cre/Kiss1^fl/fl^ mice (Nandankar et al. 2021). A tenable explanation is that *Pdyn*-Cre mediated ablation causes a nearly complete elimination of ARC Kiss1 expression, while *Tac2*-driven elimination allows discrimination between KNDy vs. non KNDy-born kisspeptins, therefore teasing apart the functions of these two compartments of kisspeptin output within the ARC.

ARC KNDy neurons and/or kisspeptin signaling have been suggested to participate in the control of bodily functions other than reproduction, including body temperature and weight regulation, as well as metabolic homeostasis (Mittelman-Smith, Williams, Krajewski-Hall, McMullen, et al. 2012; Nestor et al. 2016; Velasco et al. 2019; Mittelman-Smith, Williams, Krajewski-Hall, Lai, et al. 2012); supporting evidence for such roles comes in part from stimulation or ablation strategies of KNDy cells. However, KNDy peptides or transmitters other than kisspeptins have been proposed to play leading roles in the control of thermal vasomotor responses, i.e., NKB (Padilla et al. 2018), or feeding, i.e., glutamate (Qiu et al. 2018), although the putative contribution of kisspeptin signaling to these phenomena cannot be completely excluded based on previous experimental evidence. Our TaKKO model offers a unique tool to further interrogate the specific roles of KNDy-born kisspeptins in the control of these non-reproductive functions, especially since alterations of sex steroids levels in this model are milder than those reported in mice congenitally lacking kisspeptin signaling (Velasco et al. 2019). In fact, systematic comparison of our current dataset and previous data from Gpr54 null models highlights putative specific functions of KNDy-derived kisspeptins in the control of body weight/composition and metabolic homeostasis. Thus, consistent with previous data in Gpr54 null mice of both sexes (Tolson et al. 2014; Velasco et al. 2019), TaKKO male and female mice displayed increased adiposity and decreased lean mass; yet, the reported overall increase in body weight gain in adult female mice congenitally lacking kisspeptin signaling (Tolson et al. 2014) was not observed in TaKKO females, whose body weight was actually modestly reduced. Likewise, a decrease in locomotor activity was observed in female, but not in male TaKKO animals; suppressed locomotor activity had been previously reported in global Gpr54 KO mice (Tolson et al. 2014), and recent evidence pointed out that toxin-mediated ablation of ARC Kiss1 neurons reduced physical activity (Padilla et al. 2019). Our data conclusively demonstrate the prominent role of KNDy-born kisspeptins in the control of locomotor activity, selectively in females. Noteworthy, previous experiments showed that ARC Kiss1 silencing not only suppressed physical activity but also perturbed temperature rhythms (Padilla et al. 2019); yet, no evidence for deregulated tail or body temperature was obtained in our TaKKO mice, suggesting that such function is conducted by KNDy-neuron signals other than kisspeptins, such as NKB (Padilla et al. 2018). Likewise, KNDy-derived kisspeptins are unlikely to be involved in the anorexigenic effect caused by ARC Kiss1 neuron activation (Qiu et al. 2018), since feeding was not altered (actually, tended to be reduced) in TaKKO mice of both sexes. On the other hand, while decreased energy expenditure was previously described in global Gpr54 null mice (Tolson et al. 2014), no changes in energy expenditure were detected in TaKKO mice, although both males and females displayed decreased weight of brown adipose tissue. In addition, despite increased adiposity, glucose homeostasis was not perturbed in TaKKO mice, and in fact glucose and insulin tolerance were modestly improved in male mice with conditional ablation of *Kiss1* in *Tac2* cells. This is in contrast with the reported state of glucose intolerance described in global Gpr54 null mice (Tolson et al. 2014; Velasco et al. 2019), and highlights the distinctive function of KNDy-born kisspeptins, which may worsen per se glucose homeostasis, vs. the global impact of ablation of kisspeptin signaling, in which the state of profound hypogonadism is likely to be a major contributing factor.

In the last decades, kisspeptins, produced by discrete populations of Kiss1 neurons, have emerged as master regulators of the reproductive axis. In this context, ARC Kiss1 neurons have been shown to play a major role in the control of GnRH pulsatility and tonic gonadotropin secretion. While a prominent subset of ARC Kiss1 neurons are KNDy, they co-exist with Kiss1-only and NKB-only neurons, as well as Kiss1 neurons at other hypothalamic and extra-hypothalamic sites, which putatively engage in the central circuits governing GnRH neurosecretion, as well as other bodily functions. We present herein the first model of conditional ablation of *Kiss1* in *Tac2*-expressing neurons. Indeed, rather than global knockout of *Kiss1* in the ARC, our line allows selectively elimination of KNDy-born kisspeptins, without altering the other compartments of kisspeptin output. This represents a unique tool for in vivo dissection of the specific roles of KNDy-born kisspeptins and deepening of our knowledge of the intimate mechanisms whereby different subpopulations of Kiss1 neurons regulate various facets of the reproductive axis, and related body functions. Importantly, our data provide novel physiological evidence for the differential contribution of KNDy- vs. non KNDy-kisspeptins in the control of reproduction, and conclusively document that, while dispensable/compensable for pubertal maturation, KNDy-born kisspeptins are required for maintenance of gonadotropin pulsatility and fertility, specifically in the female. In-depth characterization of such functional diversity has become specially relevant, considering the promising potential of kisspeptin-based therapies for management of some reproductive disorders in humans (Abbara et al. 2021).

## Methods

### Ethics Statement

The experiments and animal protocols included in this study were approved by the Ethical Committee of the University of Córdoba; all experiments were conducted in accordance with European Union (EU) normative for the use and care of experimental animals (EU Directive 2010/63/UE, September 2010).

### Animals

Mice were housed in the Service of Animal Experimentation (SAEX) of the University of Córdoba. All animals were maintained in constant conditions of light, 12-h light/dark cycle, at standard temperature (22 ± 2°C), with ad libitum access to standard laboratory mice chow (A04, Panlab) and water. The day the litters were born was considered as day 1 of age (PND1); animals were weaned at PND21.

### Generation of the TaKKO mouse line

A mouse line with conditional ablation of *Kiss1* in *Tac2*-expressing cells was generated as a means to selectively eliminate kisspeptins produced by KNDy cells (which co-express *Tac2* and *Kiss1*), vs. those released by Kiss1-only neurons. To this end, the *Tac2*-IRES-Cre mouse line (B6;129S-*Tac2^tm1.1(cre)Hze^*/J; JAX:021878) was crossed with the novel Kiss1^loxP/loxP^ mouse, produced at the Turku Center for Disease Modeling (TCDM. Turku, Finland), in which exon 3 of *Kiss1* gene is flanked by two loxP sites, to allow Cre-mediated recombination. The resulting double transgenic line (Tac2-Cre::Kiss1^fl/fl^) was named TaKKO, for **<u>Ta</u>**c2 cell-specific **<u>Ki</u>**ss1 **<u>KO</u>**.

### Genotyping and validation of the TaKKO mouse line

For genotyping of TaKKO mice, PCR analyses were conducted on genomic DNA, isolated from mouse ear tissue. Primers to detect *Cre* in the *Tac2* locus were Forward: 5’-GAG ATG TGG TTC CTG GCT GT-3’; and Reverse: 5’-GGA TTG GGA AGA CAA TAG CAG G-3’, while primers used to detect the WT allele, i.e., absence of Cre, were Forward: 5’- CGA CGT GGT TGA AGA GAA CA-3’; and Reverse: 5’-GAG ATG TGG TTC CTG GCT GT-3’. In addition, detection of the loxP-flanked Kiss1 allele was conducted using the primers, Kiss1-Flox Fw: 5’- AAT GAG CAC GTA TTG GAG CC-3’; and Kiss1-Flox Plus: 5’- GCG GAT TTG TCC TAC TCA GG-3’; to detect the WT allele, the primers used were Kiss1-Flox Fw: 5’- AAT GAG CAC GTA TTG GAG CC-3’ and Kiss1- Flox Rv: 5’-TGT CTG GAG TCT GAG CCA GC-3’.

To confirm effective Cre-mediated ablation of exon 3 in the *Kiss1* gene, analysis of recombination was conducted by PCR selectively in the ARC (where KNDy neurons are located), as well as in the preoptic area (POA) and the cortex of control and TaKKO mice. The primers used for detection of recombinant band were Forward: 5’-AAT GAG CAC GTA TTG GAG CC-3’; and Reverse: 5’-TAA GCC GTT GGG TTG GAC TC-3’.

### Double immunohistochemical analysis in TaKKO mice

In order to further validate our mouse line, double immunofluorescence assays were performed to assess co-expression of kisspeptins (KP) and NKB in the ARC of control and TaKKO mice of both sexes. Mice were anesthetized with an intraperitoneal (ip) injection of ketamine/xylazine and subjected to intracardial perfusion with 0,9% saline, followed by 4% paraformaldehyde (PFA, pH 7.4) in PBS. Brains were collected and post-fixed in 4% PFA (4 °C, 24 h), washed in PBS (4 °C for 24 h), and dehydrated in sucrose (30% in 0.1 M PBS for 24–48 h). Brains were then frozen and stored at -80°C until being processed.

Twenty five-μm thick coronal sections were collected with a Leica SM 2000R freezing microtome into tissue culture plates filled with anti-freeze solution (30% ethylene glycol, 25% glycerol, 0.05 M phosphate buffer, pH 7.4) and stored at -20°C. Detection of kisspeptin- and NKB-IR with immunofluorescent double-labeling used two sequential rounds of tyramide signal amplification to maximize both signals. ARC sections from gonadectomized male and female control and TaKKO mice (n=2/each) were rinsed in PBS, pre-treated with a mixture of 1% H_2_O_2_ and 0.5% Triton X-100 for 20 min, and subjected to antigen retrieval with 0.1M citrate buffer (pH 6.0) at 80°C for 30 min. Then, a mixture of rabbit kisspeptin (AC566; 1:20.000) (Franceschini et al. 2006), and guinea pig NKB (IS-3/61; 1:20.000) (Ciofi et al. 1994) primary antibodies was applied to the sections for 48 h at 4 °C, followed by peroxidase-conjugated anti-rabbit IgG (Jackson ImmunoResearch; 1:250; 1 h) and Cy3-tyramide (diluted 1:1000 with 0.05 M Tris-HCl buffer/0.005% H2O2, pH 7.6, 30 min (Hopman et al. 1998). Peroxidase was inactivated with 1% H_2_O_2_ and 0.1 M sodium azide in PBS for 30 min. Then, the guinea pig NKB antibodies were reacted with biotin-conjugated secondary antibodies (Jackson ImmunoResearch; 1:500; 1 h), ABC Elite reagent (Vector; 1:1000; 1 h), and FITC-tyramide (diluted 1:1000 with 0.05 M Tris-HCl buffer/0.005% H2O2, pH 7.6, 30 min (Hopman et al. 1998). The dual-labeled sections were mounted and coverslipped with the aqueous mounting medium Mowiol. Fluorescent signals were studied with a Zeiss LSM780 confocal microscope. High-resolution images were captured using a 20×/0.8 NA objective, a 0.6× optical zoom, and the Zen software (Carl Zeiss). Different fluorochromes were detected with laser lines 488 nm for FITC and 561 nm for Cy3. Emission filters were 493–556 nm for FITC and 570–624 nm for Cy3. To prevent emission crosstalk between the fluorophores, the red channel was recorded separately from the green one (‘smart setup’ function). To illustrate the results, confocal Z-stacks (Z-steps: 0.974 μm; pixel dwell time: 3.15 μs; resolution: 1024×1024 pixels; pinhole size: set at 1 Airy unit) were merged using maximum intensity Z-projection (ImageJ). The final figures were adjusted in Adobe Photoshop and saved as TIF files.

### In situ hybridization analysis in TaKKO mice

Hypothalamic expression of *Kiss1* and *Tac2* mRNAs was assessed by in situ hybridization (ISH). In brief, five sets of coronal brain sections of 20-μm thick including from rostral to caudal hypothalamus were generated, mounted on Super-Frost Plus slides (Thermo Fisher Scientific, Inc.), and stored at −80 °C until ISH analyses. Specific antisense riboprobes for mouse *Kiss1* and *Tac2* mRNA, located between 82-371 nucleotides (Genbank: NM_178260.3) and 177-440 nucleotides (GenBank: NM_009312) of the cDNA sequence, respectively, were generated. For the generation of the *Kiss1* probe, the sequences of the sense and antisense primers were as follows: T3-Kiss1 Fw (5’-CAG AGA TGC AAT TAA CCC TCC TCA CTA AAG GGA GAT GGT GAA CCC TGA ACC CAC A-3’) and T7-Kiss1 Rv (5’-CCA AGC CTT CTA ATA CGA CTC ACT ATA GGG AGA ACC TGC CTC CTG CCG TAG CG-3’). For the *Tac2* probe, the sense and antisense primers used were: T3-Tac2 Fw (5’-CAG AGA TGC AAT TAA CCC TCA TCA CTA AAG GGA GAA GGG AGG GAG GAG GCT CAG TAA G-3’) and T7-Tac2 Rv (5’-CCA AGC CTT CTA ATA CGA CGA CTC ACT ATA GGG AGA TTT GAG GAG GAT GCC AAA GCT G-3’). Hypothalamic sections were fixed in 4% paraformaldehyde, stabilized with 0.1M phosphate buffer (pH 7.4) and pre-treated with triethanolamine and anhydrous acetic acid to decrease nonspecific binding. Tissues were then dehydrated in increasing ethanol concentration. Riboprobe hybridization for *Kiss1* or *Tac2* was performed for 16 hours at 55°C. The hybridization solution contained 4x SCC (saline sodium citrate sodium buffer), 50% deionized formamide, 1x Denhardt’s solution and 50% dextran sulphate. Each riboprobes was added to hybridization solution at a final concentration of 0.03 pmol/ml along with tRNA. After hybridization, slides were washed, treated with RNase-A, and dehydrated in increasing ethanol series as previously described (Manfredi-Lozano et al. 2016). Finally, slides were dipped in Kodak autoradiography emulsion type NTB (Eastman Kodak) and exposed for 2 weeks at 4 °C in the dark. After this period, the sections were developed and fixed following manufacturer instructions (Kodak). Then, slices were cover slipped with Sub-X mounting medium (Leica). For analysis, slides were read under dark-field illumination with custom-designed software and enabled to count the total number of cells (grain clusters).

### Phenotypic evaluation of pubertal maturation and reproductive assessment in TaKKO mice

After weaning, 3-week-old male and female mice were daily checked for phenotypic markers of puberty onset. Somatic and reproductive indices of pubertal development included body weight (BW) gain after weaning, age of vaginal opening (VO), as external marker of puberty onset in female rodents, and balano-preputial separation (BPS), as sign of pubertal maturation in male rodents. Once the VO occurred, vaginal cytology was conducted daily to identify the age of the first estrus (FE), as phenotypic sign of first ovulation in female rodents. Adult virgin female mice, at 2- and 12-months of age, were monitored daily by vaginal cytology for at least 2-3 weeks to characterize the estrous cyclicity. Histological analyses were conducted in ovarian samples from 4-week, 2-month and 12-month-old female mice, and in testicular samples from adult (10-12 month-old) males.

### Basal hormonal analyses in TaKKO mice

Basal levels of LH and FSH were measured in single time-point serum samples, using RIA kits supplied by the National Institutes of Health (Dr. A. F. Parlow, National Hormone and Peptide Program, Torrance, CA). Rat LH-I-10 and FSH-I-9 were labelled with ^125^I by the chloramine-T method. LH and FSH concentrations were expressed using reference preparations LH-RP-3 and FSH-RP-2, respectively, as standards. Intra and inter-assay coefficients of variation were <8% and <10% for LH and <6% and <9% for FSH. The sensitivity of the assay was 5 pg/tube for LH and 20 pg/tube for FSH. In addition, sex steroid levels were measured in serum samples using a previously validated, sensitive liquid chromatography-tandem mass spectrometry (LC-MS/MS) method, adapted from protocols described in detail elsewhere (Luque-Cordoba and Priego-Capote 2021). Determinations of estradiol (E_2_), testosterone (T), dihydro-testosterone (DHT) and progesterone (P) were conducted in control and TaKKO mice of both sexes, at 4-weeks and 2-months of age.

### Assessment of pulsatile and surge LH secretion in TaKKO mice

For analysis of pulsatile secretion of LH, intact female mice were handled (5-10 min per animal) every day for 3 weeks before blood sampling, in order to train them for tail-tip bleeding. For blood sample collection, diestrus female mice were restrained in a cardboard tube and 4µl of blood were obtained from the mouse-tail every 5 minutes interval during 3-hours. Blood samples were diluted in 46 μl of 0.1 M PBS (phosphate buffered saline) with 0.05 % Tween 20, snap-frozen on dry ice, and finally samples were stored at -80°C until assayed using a super-sensitive LH ELISA (Steyn et al. 2013). The basal and mean LH levels, as well as the number of pulses were measured. In this case, LH pulses were defined as those displaying an increase of 125% over the 5 lowest points, based on validated protocols (Franssen et al. 2021; Torres et al. 2021), adapted from previous references (McQuillan et al. 2019). Basal LH levels were calculated using the mean of the values preceding each peak, whereas mean LH levels were estimated as the arithmetic mean of all LH values over the study period. Pulse amplitude was calculated as the difference between the pulse peak and the preceding point, and area under the curve (AUC) were estimated by the trapezoidal rule method.

In addition, sex steroid-priming was applied to evoke LH preovulatory peaks, analyzed in female mice subjected to bilateral ovariectomy (OVX), via abdominal incision under light isoflurane anesthesia, followed by replacement with physiological concentrations of estrogen. The replacement consisted of a subcutaneous implant of a SILASTIC capsule (Dow Corning, Milland, MI, USA), whose characteristics were as follows: total length of 1.5 cm; effective length of 1 cm; internal diameter of 1.98 mm and external diameter of 3.18 mm. Animals were diary checked for vaginal smears and once all the animals had changed to diestrus, approximately at day 9 after OVX+E_2_ replacement, they were injected (10:00-11:00 am) with estradiol benzoate (1µg per 20g of body weight). The day after, basal blood samples from mouse-tail were collected at 10:00am and 7:00pm (light phase) and, immediately after lights off, blood samples were collected at 3 additional time-points every 15 minutes, as adapted from previously published protocols (Leon et al. 2021; Czieselsky et al. 2016). Assessment of LH levels in these serial samples was conducted by a super-sensitive ELISA, as described above.

### Fertility tests and gonadal histology in TaKKO mice

To assess fertility, individual breeding pairs were monitored during a 3-month period, allowing to record a minimum of 3 breeding cycles. Animals were paired as follow: control male with control female (n=5); TaKKO male with control female (n=6); and control male with TaKKO female (n=5). Mice were checked daily for number of pups and the day of birth of each litter was recorded. After birth, the pups were immediately euthanized, and the breeding pair was followed for the remainder 3-month.

In addition, histological analyses were conducted in male and female gonads from control and TaKKO mice. As general procedure for tissue processing, after collection at euthanasia, testes and ovaries, together with the oviduct and the tip of the uterine horn, were fixed for at least 24 hours in Bouin solution and processed for paraffin embedding. The gonads were serially sectioned (10 μm-thick), stained with hematoxylin and eosin, and viewed under an Elipse 400 microscope for evaluation. For ovarian studies, qualitative assessment of the presences of follicles, at different maturational stages, as well as the presence and number of corpora lutea, taken an index of ovulation, were recorded (Gaytan et al. 2017). For testicular studies, histological analyses included qualitative assessment of the general structure of testicular compartments and the different stages spermatogenesis, in line with a previous reference (Endo et al. 2017).

### Basic metabolic phenotyping in TaKKO mice

Body composition, including fat mass and lean mass, was measured by quantitative magnetic resonance imaging with an EchoMRI™ 700 analyzer (Houston, TX, software v.2.0). In addition, mice were placed into metabolic chambers to analyse different metabolic parameters such as energy expenditure (EE), respiratory quotient (RQ) and locomotor activity. Oxilet pro: Intake & Activity (LE1335), O_2_/CO_2_ Analyzer (LE405) and Air switch and Flow checking (LE4004FL), Harvard Apparatus serial number 1791914, were used to determine EE and RQ. Locomotor activity, based on the monitoring of the movement of the animal in a horizontal plane, was recorded every 3 minutes.

### Glucose and insulin tolerance tests in TaKKO mice

To assess glucose homeostasis, adult female and male mice were subjected to glucose tolerance tests (GTT). Mice (n=9-11 animals/group) were fasted for 4h and subsequently received an ip bolus of glucose (1,5 g/kg BW). Glucose levels were determined before (0-min) and at 20, 60, and 120-min post administration. Glucose concentrations were measured using a handheld glucometer (ACCU-CHECK Aviva; Roche Diagnostics). Integral glucose changes were estimated by the trapezoidal rule method as AUC during the 120 min period. In addition, insulin tolerance tests (ITT) were performed in adult female and male mice. Mice (n=9-11 animals/group) were fasted 4h and glucose levels were measured before (0-min) and at 20, 60, and 120-min after ip injection of insulin (0,75UI). Integral glucose changes were estimated by trapezoidal method as AUC during the 120 min following insulin administration.

### Temperature analyses in TaKKO mice

To assess heat dissipation by vasodilation in the tail, an ultra-small temperature logger (DST, Nano-T, Star-Oddi, Garðabær, Iceland) was attached to the ventral surface at the base of the tail. The mice were individually caged allowing free movement and ad libitum access to water and food. Tail temperatures were recorded every 10 minutes for 24h with a previous 1-day period of habituation. In addition, interscapular back temperature was measured and visualized using a high-resolution infrared camera (E54 Compact-Infrared-Thermal-Imaging-Camera; FLIR systems OÜ, Estonia) and analysed with a FLIR-Tools specific software package. Animals were caged individually and interscapular zone was shaved prior to BAT/back temperature measures.

### Statistical analyses

Statistical analyses were performed using Prism software (GraphPad Prism). All data are presented as mean ± standard error of the mean (SEM). Group sizes (defined as biological replicates) for each experiment are indicated in the figure legends. Sample sizes were defined based on previous experience of the group in experimental studies using rodents, addressing to evaluate neuroendocrine regulation of puberty and adult reproductive function, and were assisted by power analyses performed using values of standard deviation that we usually obtain when measuring parameters similar to those examined herein. The selected sample sizes were predicted to provide at least 80% power to detect effect sizes using the tests indicated above, with a significance level of 0.05. Nonetheless, according to standard procedures, more complex molecular and histological analyses were implemented in a representative subset of randomly assigned samples from each group. Unpaired t-tests were applied for assessment of differences between two groups, while ANOVA followed by post hoc multiple comparison Tukey tests for comparisons of more than two groups. The significance level was set at P ≤ 0.05. As general rule, the investigators directly performing animal experimentation and analyses were not blinded to group allocation, but primary data analyses by senior authors were conducted independently to avoid any potential bias.

### Data availability

The authors declare that the data supporting the findings of this study are included in this article and its supplementary information files. All relevant original data are provided also as source data files, included as supplementary material.

## Acknowledgements

This work was supported by grants BFU2017-83934-P and PID2020-118660GB-I00 (Agencia Estatal de Investigación, Spain; co-funded with EU funds from FEDER Program); grant NNF18OC0034370 (Novo Nordisk Foundation); FiDiPro (*Finnish Distinguished Professor*) Program of the Academy of Finland (to M.P. and M.T.-S.); project PIE14-00005 (Flexi-Met, Instituto de Salud Carlos III, Ministerio de Sanidad, Spain); Projects P12-FQM-01943 and P18-RT-4093 (Junta de Andalucía, Spain), Project 1254821 (University of Cordoba-FEDER); National Science Foundation of Hungary (K128317, K138137, PD134837; to E.H.); and EU research contract GAP-2014-655232. CIBER is an initiative of Instituto de Salud Carlos III (Ministerio de Sanidad, Spain).

---

**Suppl. Table S1:**
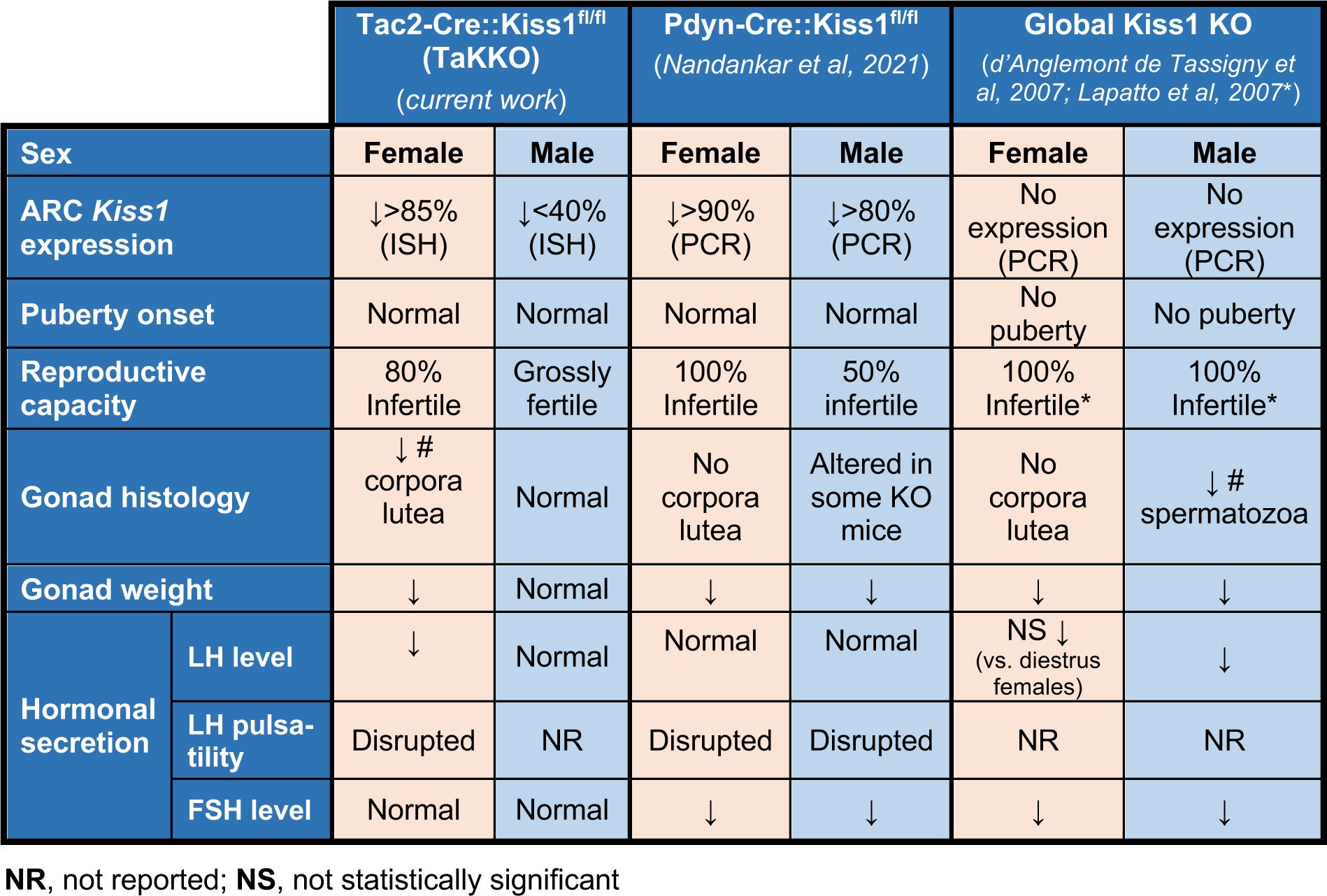
Comparative phenotypes of Tac2-specific Kiss1 null (TaKKO) mice vs. other mouse models of conditional or global ablation of Kiss1. Basic molecular, phenotypic and hormonal features of TaKKO mice are compared with those of a model of conditional ablation of Kiss1 in Pdyn-expressing cells (Pdyn-Cre::Kiss1^fl/fl^) and those of mouse lines with global conditional ablation of Kiss1. References are indicated.

|  |  | **Tac2-Cre::Kiss1<sup>fl/fl</sup>**<br>**(TaKKO)**<br>(current work) |  | **Pdyn-Cre::Kiss1<sup>fl/fl</sup>**<br>(Nandankar et al, 2021) |  | **Global Kiss1 KO**<br>(d'Anglemont de Tassigny et al, 2007; Lapatto et al, 2007*) |  |
| --- | --- | --- | --- | --- | --- | --- | --- |
| **Sex** |  | **Female** | **Male** | **Female** | **Male** | **Female** | **Male** |
| **ARC *Kiss1* expression** |  | ↓>85% (ISH) | ↓<40% (ISH) | ↓>90% (PCR) | ↓>80% (PCR) | No expression (PCR) | No expression (PCR) |
| **Puberty onset** |  | Normal | Normal | Normal | Normal | No puberty | No puberty |
| **Reproductive capacity** |  | 80% Infertile | Grossly fertile | 100% Infertile | 50% infertile | 100% Infertile* | 100% Infertile* |
| **Gonad histology** |  | ↓ # corpora lutea | Normal | No corpora lutea | Altered in some KO mice | No corpora lutea | ↓ # spermatozoa |
| **Gonad weight** |  | ↓ | Normal | ↓ | ↓ | ↓ | ↓ |
| **Hormonal secretion** | **LH level** | ↓ | Normal | Normal | Normal | NS ↓ (vs. diestrus females) | ↓ |
|  | **LH pulsatility** | Disrupted | NR | Disrupted | Disrupted | NR | NR |
|  | **FSH level** | Normal | Normal | ↓ | ↓ | ↓ | ↓ |

**NR**, not reported; **NS**, not statistically significant

## Legend to Supplementary Figures

**Suppl. Figure S1:**
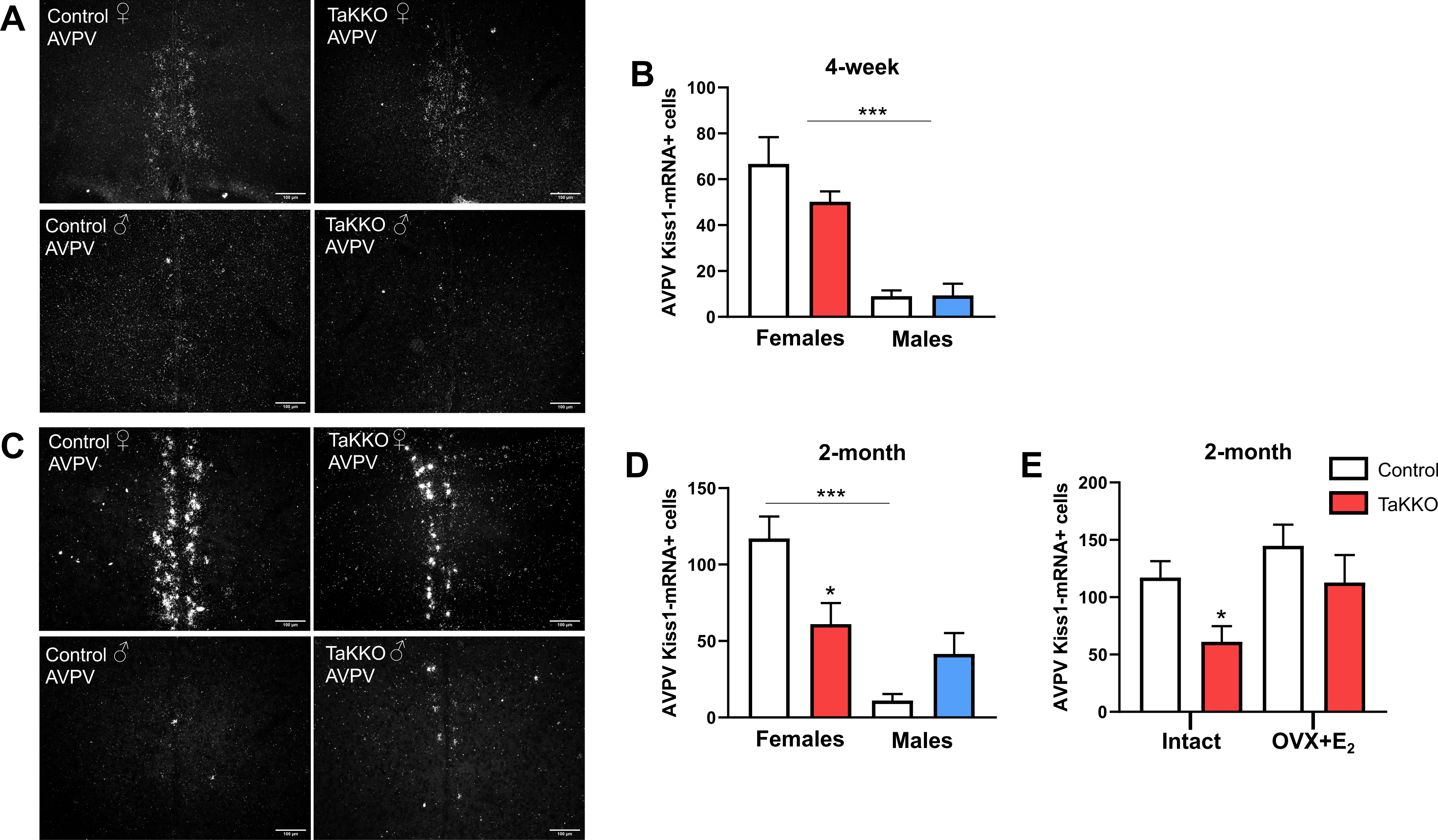
Changes in Kiss1 expression in the AVPV of TaKKO mice during postnatal maturation. Representative ISH photomicrographs are shown of *Kiss1* mRNA expression in the AVPV, from 4-week (peri-pubertal; panel **A**) and 2-month-old (young-adult; panel **C**) control and TaKKO female and male mice. *Kiss1* mRNA expression was quantified as estimated by grain density in the areas subjected to analysis in 4-week (panel **B**) and 2-month-old (panel **D**) in female and male mice. In addition, in panel **E**, quantification of *Kiss1* mRNA expression in the AVPV of intact and ovariectomized female mice of both genotypes, subjected to physiological replacement with estradiol (E_2_), is shown. Data are presented as mean ± SEM. Statistical significance was determined by was determined by ANOVA followed by Tukey’s post hoc analysis: *P<0.05 vs. corresponding control group; ^+++^ P<0.001 vs. opposite sex of the same genotype. The scale bar represents 100µm.

**Suppl. Figure S2:**
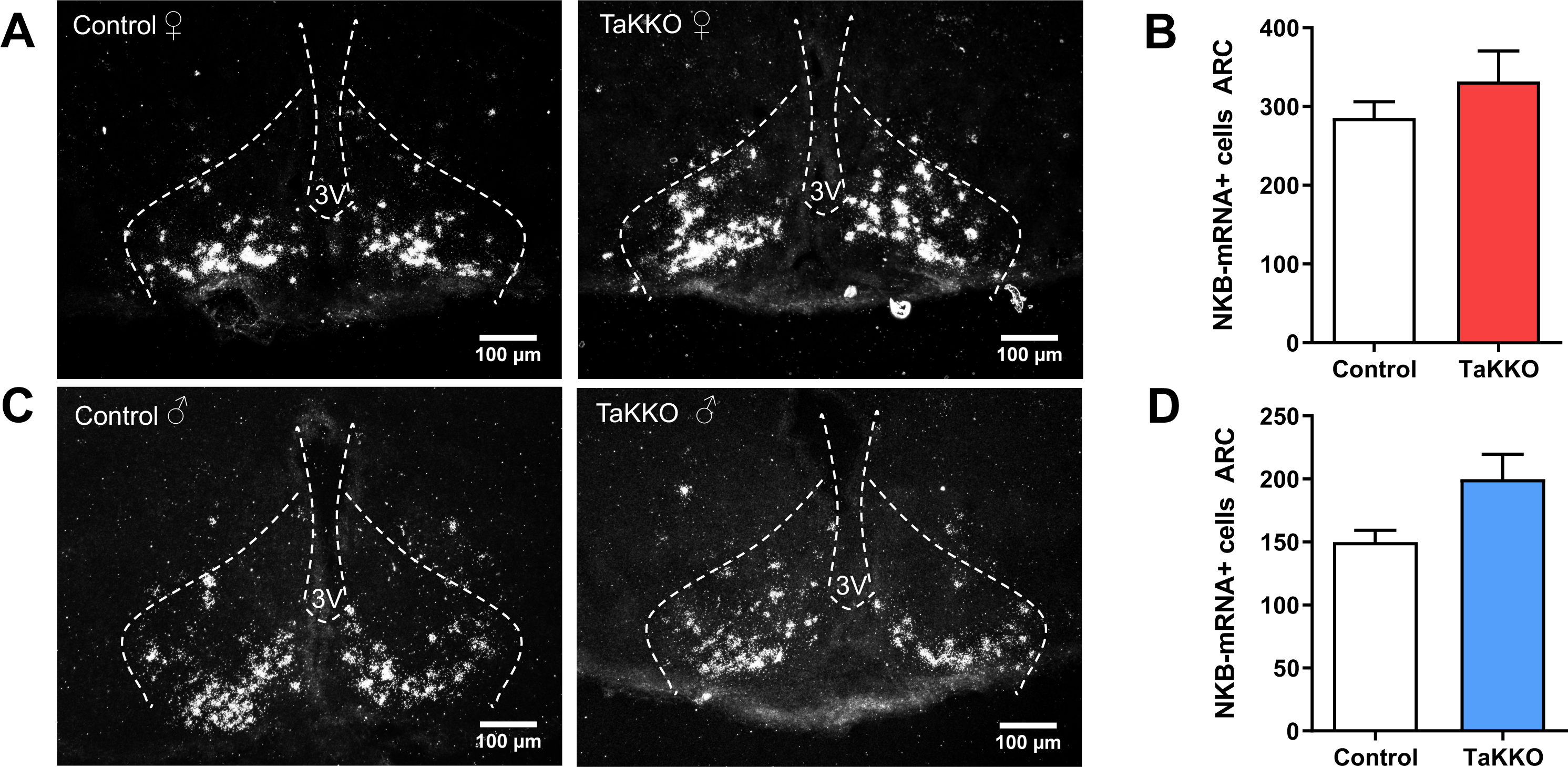
Tac2 expression in the ARC of TaKKO mice. In panel **A**, representative ISH photomicrographs are shown of *Tac2* (encoding NKB)-expressing neurons in the anterior region of the ARC in young adult (2-mo-old) control and TaKKO female mice (Control n= 5; TaKKO n= 5). In panel **C**, representative ISH photomicrographs of *Tac2*-expressing neurons in the anterior region of the ARC at young adult control and TaKKO male mice are shown (Control n= 5; TaKKO n= 5). In addition, quantification of NKB (as *Tac2*-expressing) cells, as estimated by grain density and clusters in the areas subjected to analysis, in female (panel **B**) and male (panel **D**) mice is shown. Data are presented as mean ± SEM. No statistically significant differences were detected between the groups (Student t-test). The scale bar represents 100µm.

**Suppl. Figure S3:**
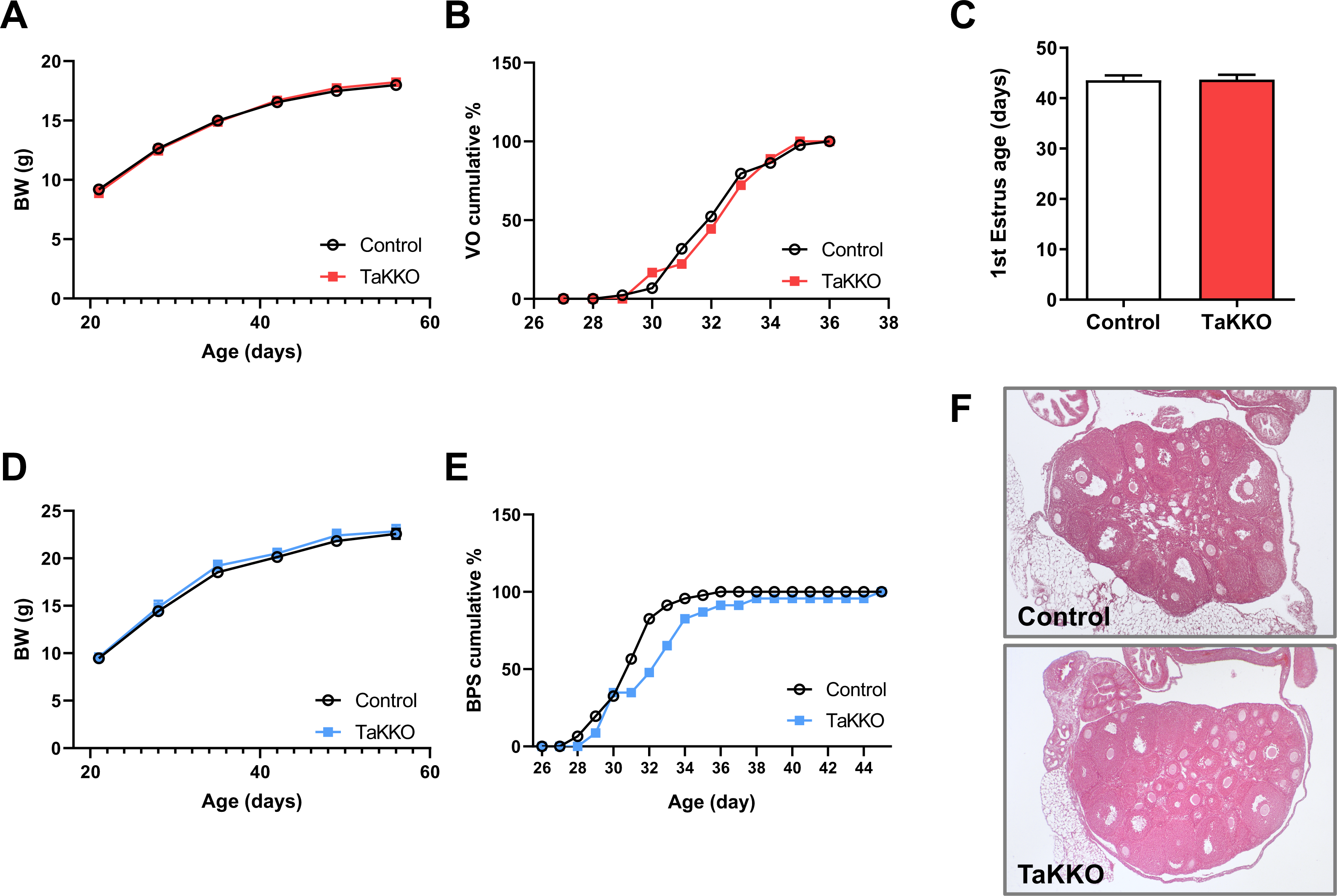
Assessment of pubertal maturation in TaKKO mice. In panel **A**, evolution of body weight (BW) gain from weaning (PND21) to adulthood (PND56) is presented in control and TaKKO female mice (Control n=23; TaKKO n=15). In panel **B**, the cumulative percentage of vaginal opening (VO) in female mice is shown, while the mean age of first estrus, as sign of first ovulation, is presented in panel **C** (Control=44; TaKKO=18). In addition, in panel **D**, we present the evolution of BW gain from weaning (PND21) to adulthood (PND56) in control and TaKKO male mice (Control n=23; TaKKO n=8). In panel **E**, the cumulative percentage of male mice displaying balano-preputial separation (BPS), as external sign of male puberty, is shown (Control=46; TaKKO=23). Finally, in panel **F**, representative images of ovarian maturation in control and TaKKO female mice at PND28, corresponding with early pubertal period, preceding the first ovulation, are shown. Note that large healthy antral follicles are present in both genotypes.

**Suppl. Figure S4:**
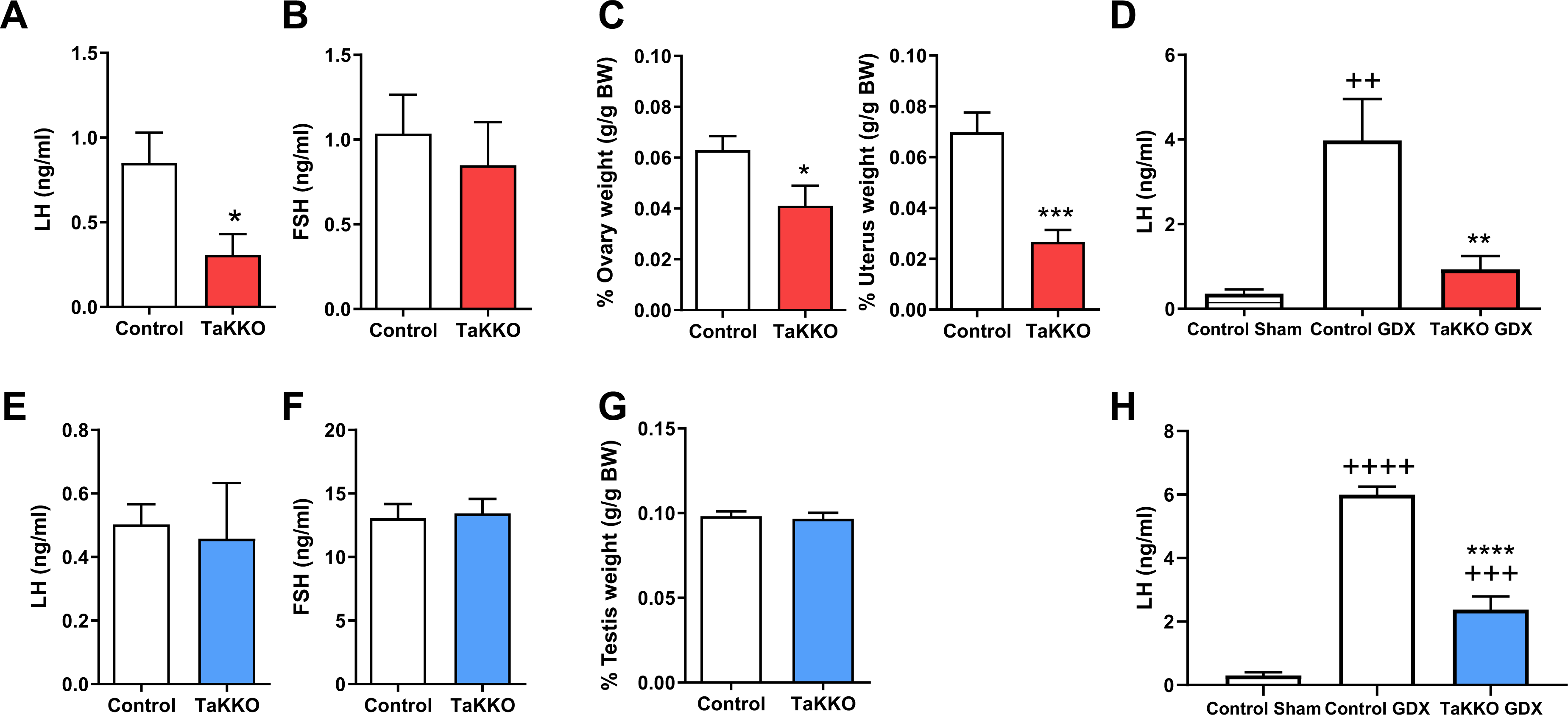
Basic reproductive parameters of young adult TaKKO mice. Basal gonadotropin levels, as well as LH responses to gonadectomy (GNX) and sex organ weights are presented from male and female TaKKO mice and their controls, at 2-months of age. Basal levels of LH and FSH are presented of female (panels **A-B**) and male (panel **E-F**) control and TaKKO mice. In addition, in panel **C**, relative ovarian and uterus weights in young adult control and TaKKO female mice are shown, while in panel **G**, relative testis weights are presented from control and TaKKO male mice. Finally, LH responses to gonadectomy (GNX) in female and male control and TaKKO are displayed in panels **D** (females) and **H** (males). Data are presented as mean ± SEM. Statistical significance was determined by Student t-test (panels A-C; E-G): *P<0.05, and ***P<0.001 vs. controls; or ANOVA follow by multiple comparison Tukey tests (panels D, H): ++P<0.005, +++P<0,0005 and ++++P<0.0001 vs control-sham; and **P<0.005 and ****P< 0.0001 vs control-GDX.

**Suppl. Figure S5:**
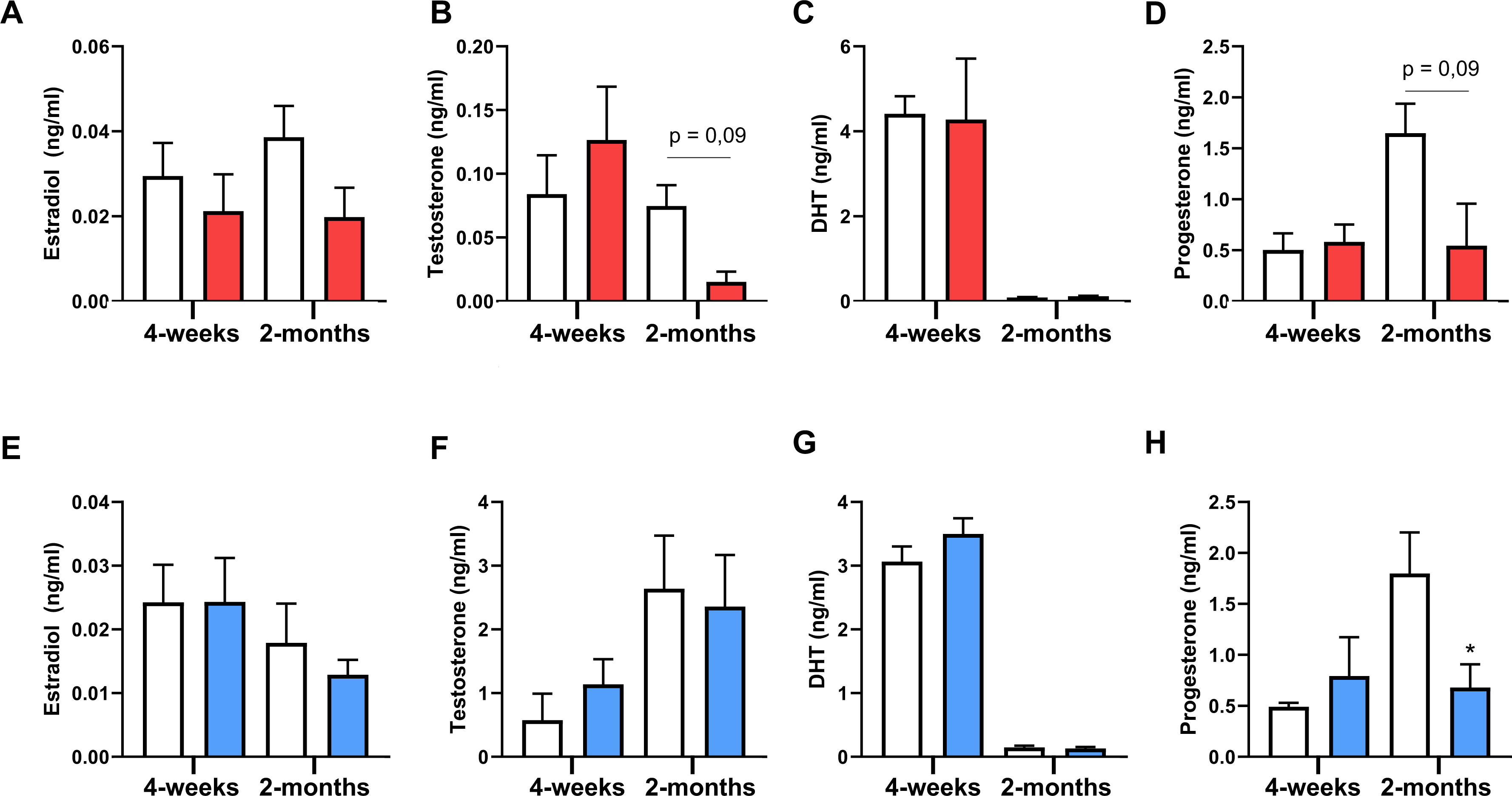
Sex steroid levels in TaKKO mice of both sexes. Histograms represents serum levels of estradiol (panels **A**, **E**), testosterone (panels **B**, **F**), dihydrotestosterone (DHT; panels **C**, **G**) and progesterone (panels **D**, **H**) in control and TaKKO female (*upper panels*) and male (*lower panels*) mice, at 4-weeks and 2-month-old mice. Data are presented as mean ± SEM. Due to large variability between the two ages, statistical significance was determined between genotypes within each age-point, using Student t-test: *P<0.05 vs. corresponding controls.

**Suppl. Figure S6:**
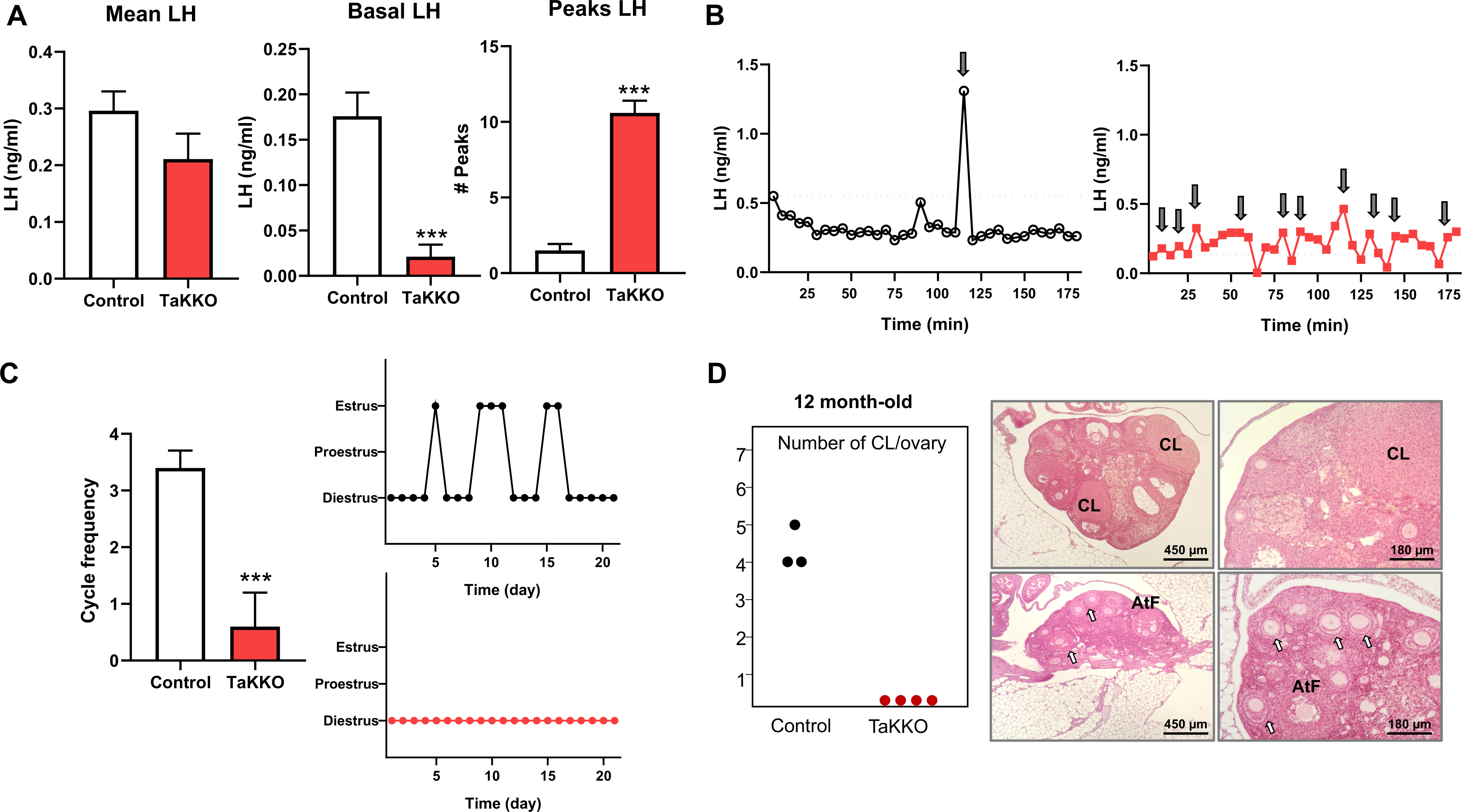
Reproductive and fertility parameters of 12-mo-old TaKKO female mice. In panel **A**, mean and basal LH levels, as well as number of LH pulses, detected over a 3-hour sampling period, are represented from aged (12-mo-old) control and TaKKO female mice (Control n=6 and TaKKO n=5). In panel **B**, individual representative LH secretory profiles of control (black line) and TaKKO (in red) female mice are shown. In addition, in panel **C**, indices of estrous cyclicity, including frequency of cycles and representative cyclic profiles, are shown from 12-mo-old control and TaKKO female mice. Finally, in panel **D**, ovulatory efficiency, denoted by the number of corpora lutea per ovary, and representative histological images of ovarian sections from aged control and TaKKO female mice (Control n=5; TaKKO n=5) are shown. In controls, corporal lutea were present, denoting persistent ovulation, while none of the 12-mo-old TaKKO females showed corpora lutea, with large atretic follicles and apparently normal small follicles (denoted by arrows). Data are presented as mean ± SEM. Statistical significance was determined by Student t-test: ***P<0.001 vs. controls. Black arrows represent a LH pulse, considered as an increment of 125% over the five lowest basal LH values. Atretic follicles: AtF; CL: corpus luteum.

**Suppl. Figure S7:**
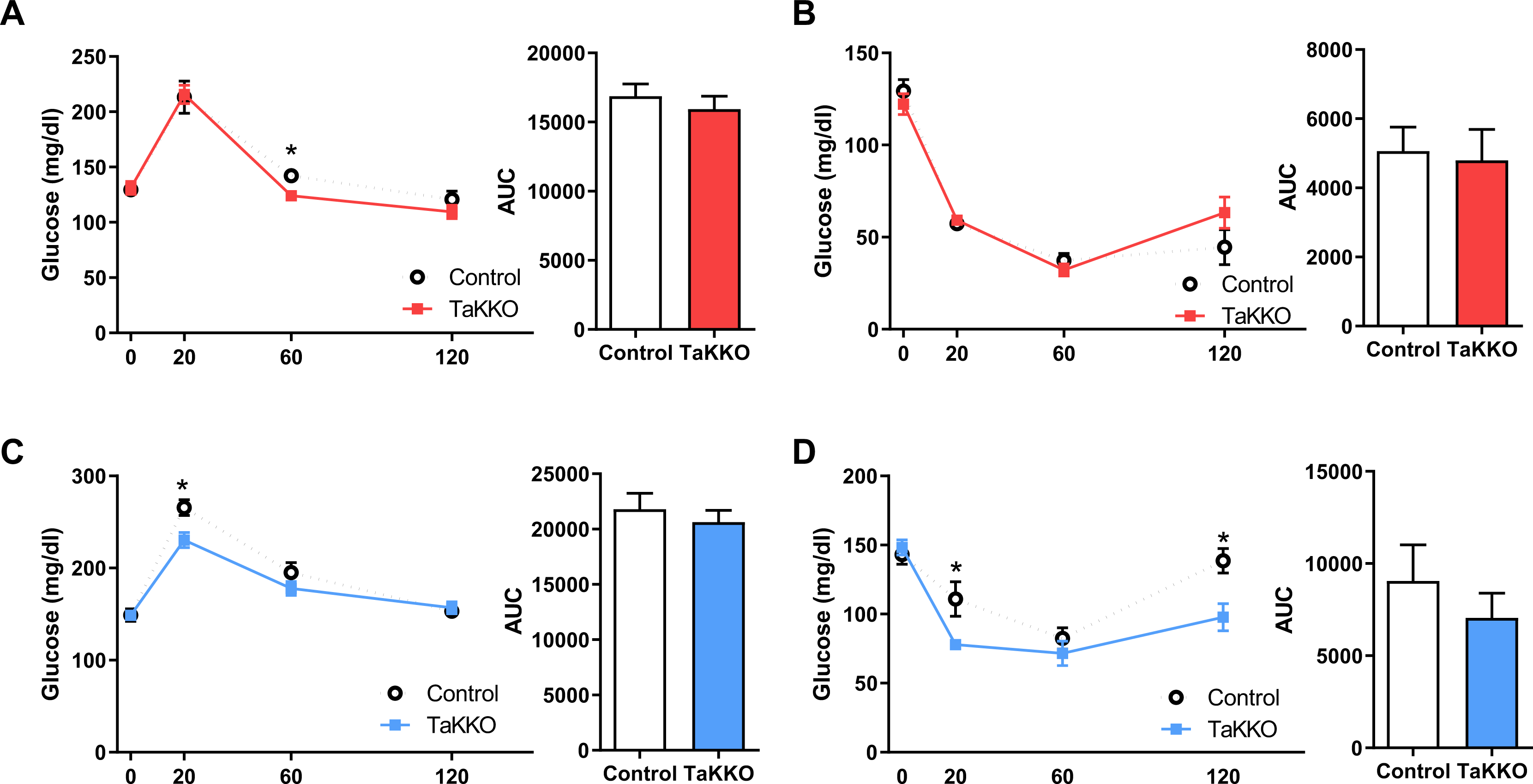
Glucose homeostasis in TaKKO mice of both sexes. In panel **A**, glucose tolerance tests (GTT) were applied to female (panel **A**) and male (panel **C**) control and TaKKO mice; values are presented as glycemic profiles and AUC. In addition, insulin tolerance tests (ITT) were conducted in female (panel **B**) and male (panel **D**) mice of both genotypes (Female: Control n=11, TaKKO n=8; male Control n=9, TaKKO n=10). Data are presented as mean ± SEM. For given time-points, statistical significance was determined by comparison between genotypes using Student t-test: *P<0.05 vs controls.

**Suppl. Figure S8:**
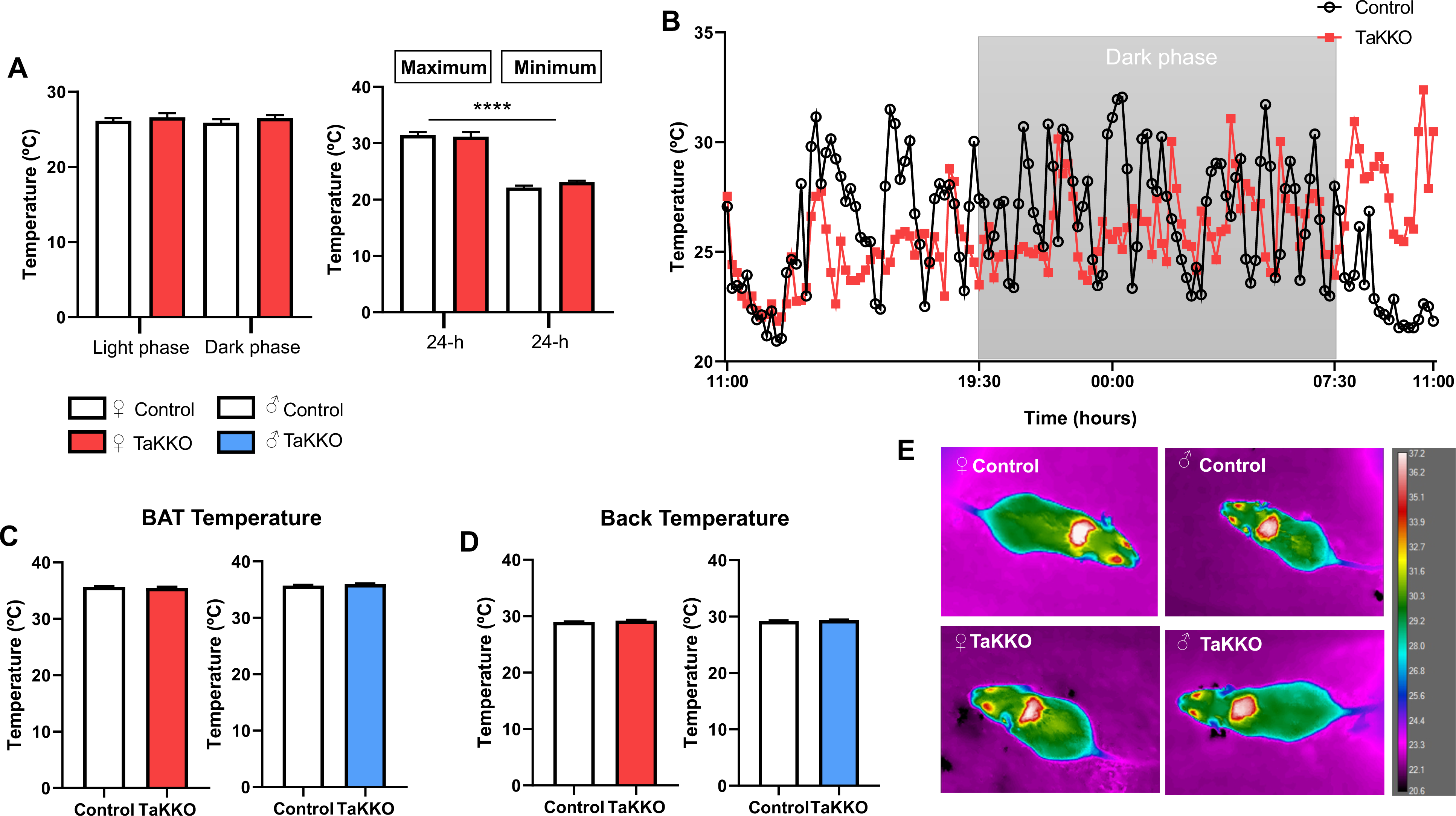
Temperature analyses in TaKKO mice of both sexes. In panel **A**, mean tail skin temperature during light and dark phases, together with maximum and minimum temperatures in a 24-h period, are recorded from control and TaKKO female mice, using a small data-logger attached to the surface of the tail. In panel **B**, individual representative temperature profiles of control (black line) and TaKKO (colour line) female mice are shown. In addition, infrared thermal quantification of the dorsal area is presented. Histograms represent BAT interscapular mean temperature (panel **C**) and back temperature (panel **D**) in control and TaKKO mice of both sexes. In addition, in panel **E**, representative thermal images in control and TaKKO female and male mice are shown. When relevant, data are presented as mean ± SEM. Statistical significance was determined by ANOVA followed by Tukey’s post hoc analysis: **P<0.01 vs. light-phase groups.

## Notes

### Competing Interest Statement

The authors have declared no competing interest.

